# Developing an evolutionary baseline model for humans: jointly inferring purifying selection with population history

**DOI:** 10.1101/2023.04.11.536488

**Authors:** Parul Johri, Susanne P. Pfeifer, Jeffrey D. Jensen

## Abstract

Building evolutionarily appropriate baseline models for natural populations is not only important for answering fundamental questions in population genetics – including quantifying the relative contributions of adaptive vs. non-adaptive processes – but it is also essential for identifying candidate loci experiencing relatively rare and episodic forms of selection (*e.g.,* positive or balancing selection). Here, a baseline model was developed for a human population of West African ancestry, the Yoruba, comprising processes constantly operating on the genome (*i.e.*, purifying and background selection, population size changes, recombination rate heterogeneity, and gene conversion). Specifically, to perform joint inference of selective effects with demography, an approximate Bayesian approach was employed that utilizes the decay of background selection effects around functional elements, taking into account genomic architecture. This approach inferred a recent 6-fold population growth together with a distribution of fitness effects that is skewed towards effectively neutral mutations. Importantly, these results further suggest that, while strong and/or frequent recurrent positive selection is inconsistent with observed data, weak to moderate positive selection is consistent but unidentifiable if rare.

## INTRODUCTION

Quantifying the relative contributions of adaptive vs. non-adaptive processes in shaping observed levels of genomic variation remains difficult. This is largely due to the fact that multiple evolutionary processes can affect patterns of variation in a similar manner, making it challenging to disentangle their individual contributions. For instance, while genetic hitchhiking effects resulting from both recurrent selective sweeps (Maynard Smith and Haigh 1974) and background selection (BGS; Charlesworth et al. 1993) may skew the allele frequency distribution towards rare alleles (Kim 2006; Nicolaisen and Desai 2012; 2013; Ewing and Jensen 2016; Johri et al. 2021), neutral population growth can result in a similar skew (see review of Charlesworth and Jensen 2021). In addition to conflicting signatures created by different evolutionary processes, heterogeneity in the rates of mutation and recombination as well as gene density across the genome add to the noise generated by these processes. Thus, in order to accurately quantify the frequency of, and identify candidate loci experiencing, rare and episodic forms of selection (such as positive selection), one must first construct an evolutionary baseline model that includes the effects of constantly acting evolutionary processes, such as genetic drift resulting from the underlying non-equilibrium population history as well as purifying and background selection caused by the constant input of deleterious mutations (Johri et al. 2022a). As most new fitness-impacting mutations are indeed deleterious (see review of Bank et al. 2014) it is particularly important to correct for them when predicting patterns of genomic variation in and around functional regions – however, the interplay of these purifying and background selection effects with population history is non-trivial (Johri et al. 2020; 2021).

Building an appropriate baseline model thus requires the quantification of parameters describing the population history as well as those defining the distribution of fitness effects (DFE) of new deleterious mutations. As accurately inferring parameters of the DFE requires corrections for the demographic history of a population (Eyre-Walker and Keightley 2007; Boyko et al. 2008), a common workaround is to follow a two-step approach whereby alleles at putatively neutral sites are utilized to obtain the demographic history, and the DFE is then inferred from variation at functional sites conditional on that estimate of demography (see review of Johri et al. 2022b). However, apart from the difficulty of identifying genuinely neutral sites in many organisms, even if these sites are successfully identified they may still experience background selection effects due to linkage with directly selected sites. As neutral demographic estimators do not account for this effect, the resulting skew in the site frequency spectrum (SFS) owing to BGS will often be misinterpreted as population growth (Ewing and Jensen 2014; Johri et al. 2021). Consequently, it is preferable to simultaneously account for the linked effects of purifying selection when inferring parameters of population history, highlighting the importance of performing joint inference of the DFE with demography.

In this study, we utilized the joint inference approach of Johri et al. (2020) in an approximate Bayesian computational framework (ABC; Beaumont et al. 2002), in order to uniquely infer the joint parameters of purifying selection and demography in a human population, the Yoruba from Ibadan, Nigeria (YRI). This approach utilizes the decay of background selection effects around functional regions while correcting for the specific genome architecture as well as the underlying heterogeneity in recombination and gene conversion rates across the genome, and has previously been shown to perform well across arbitrary DFE shapes (Johri et al. 2020; 2021). Furthermore, as the method makes no *a priori* assumptions about the neutrality of specific site types (*e.g.,* synonymous sites), it is also robust to the presence of weak selection at these sites (Johri et al. 2020). Our inference procedure suggests recent population growth, together with a DFE strongly skewed towards effectively neutral and weakly deleterious mutations. We compare this finding with previous estimates based upon two-step inference approaches, and investigate the statistical identifiability of positively selected mutations within the context of this estimated baseline model.

## RESULTS & DISCUSSION

The expected pattern of decay of background selection effects around exonic regions (see Johri et al. 2020) was used to perform the joint inference of DFE-shape with population size change in the Yoruba population, while correcting for region-specific rates of crossing over and genetic architecture. As gene conversion can significantly affect hitchhiking effects around functional genomic elements (Figure S1), region-specific rates of gene conversion were also newly incorporated into this inference framework. As the direct and linked effects of purifying selection were modeled specifically for a single exon, this method is applicable to the subset of exons in the genome for which interference effects from other nearby functional regions are minimal.

### Selecting exons in the human genome

In order to identify such exons, the recovery of nucleotide diversity (*Π*) at neutral sites due to BGS was predicted theoretically for each exon in the human genome (based upon the DFE inferred by Keightley and Eyre-Walker 2007), using equations 3a and 3b of Johri et al. (2020). More specifically, it has been shown previously (Johri et al. 2020) that if the distribution of fitness effects of new mutations follows a uniform distribution, analytical expressions for the nucleotide diversity at linked neutral sites near a functional element can be obtained. Thus, for the purpose of this study we assumed that the DFE of new deleterious mutations was comprised of four non-overlapping uniform distributions (Figure 1a), such that an *f*_0_proportion of all new mutations was neutral (2*N_e_s* = 0), an *f*_1_ proportion was weakly deleterious (1 < 2*N_e_s* ≤ 10), an *f*_2_ proportion was moderately deleterious (10 < 2*N_e_s* ≤ 100), and an *f*_3_ proportion was strongly deleterious (100 < 2*N_e_s* ≤ 2*N_e_s*), where *N_e_* is the effective population size and *s* > 0 represents the selection coefficient against homozygotes. Nucleotide diversity relative to that expected under strict neutrality, at a site that is physically at a distance *z* from a selected site, is given by:

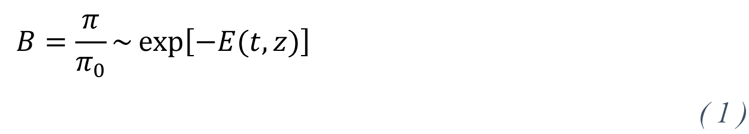

**Figure 1:**
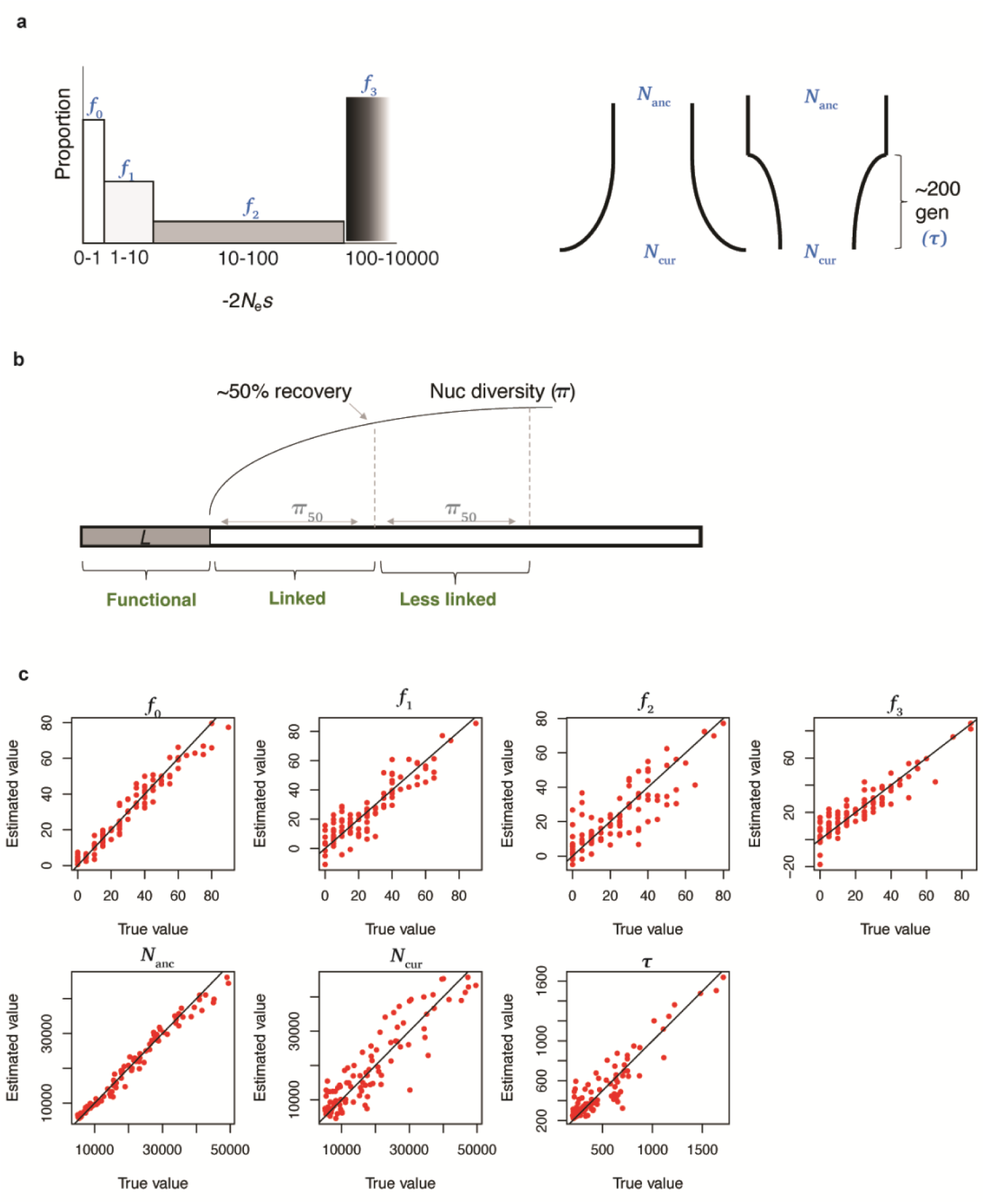
(a) Model and parameters inferred by the ABC method. The left panel shows the DFE while the right panel shows the single, recent size change demographic model fit to the data. All inferred parameters are indicated in blue font. (b) Schematic of the expected number of bases (*Π*_50_) to reach a 50% recovery of nucleotide diversity due to BGS around single exons. The three windows in which statistics were calculated are shown in green font. (c) Accuracy of joint inference of demography and the DFE. Cross-validation was performed on 100 randomly selected parameter combinations for all size parameters with tolerance = 0.08. The black line represents the *y*=*x* line. All statistics were used for inference and were calculated after removing sites inaccessible to next-generation sequencing in both the simulated and empirical data.

such that *t* = *sh* where *h* is the dominance coefficient and

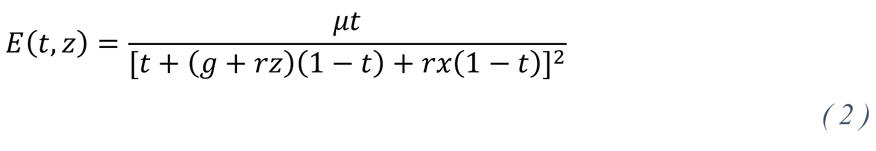

where *u* is the mutation rate, *g* is the gene conversion rate and *r* is the cross-over rate per site per generation. In order to account for BGS effects generated by a functional element of length *L*, with *t* following the probability density function *φ*(t), the expression above can be integrated over both:

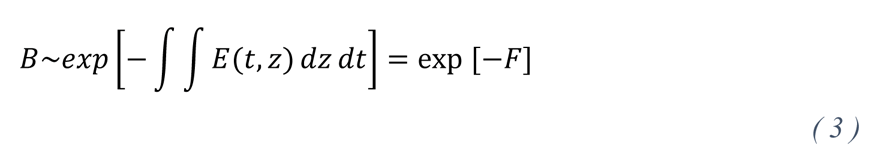

Upon integration, *F* was obtained (as shown in Johri et al. 2020) as:

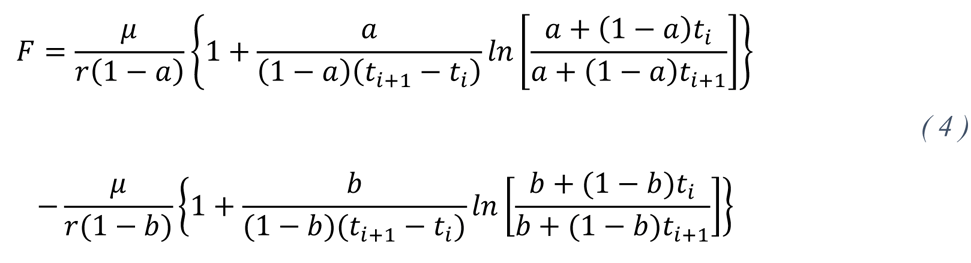

where *a* = *g* + *ry* and *b* = *g* + *r(y + L)*, where *y* is the number of sites between the neutral site and the end of the functional region and *t_i_* correspond to the boundaries of the bins. As a DFE comprised of four bins was assumed, the effect of all bins were summed up as follows:

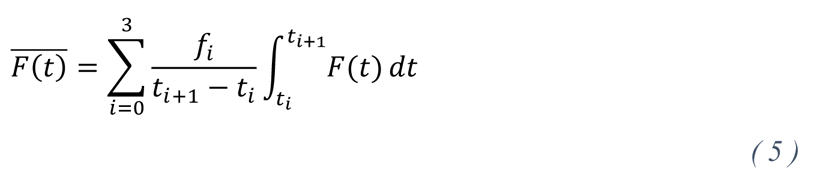

For the purpose of these analytical predictions, the gene conversion rate was assumed to be zero, which results in conservative estimates of *B*. The DFE inferred by Keightley and Eyre-Walker (2007) was assumed such that *f*_0_ = 0.22, *f*_1_ = 0.27, *f*_2_ = 0.13, *f*_3_ = 0.38 and all mutations were assumed to be semidominant. Using the above equations (1-5) derived in Johri et al. (2020), it is possible to analytically calculate expected values of nucleotide diversity as one moves away from a functional region. The expected number of bases required for a 50% recovery of diversity (*Π*_50_) was calculated as detailed in the Methods. Note that this decay of nucleotide diversity due to BGS is dependent on the length of each exon as well as the local recombination rate, and thus is specific to the human population under consideration. This analytical approach was applied to identify a subset of exons for which there were no other exons or large (> 500 bp) functionally important regions (sno/miRNAs and phastCons elements; Siepel et al. 2005) present within 4 × *Π*_50_bases of the ends of the exons. In addition, in order to observe sufficient BGS effects, our application was limited to larger exons, sized between 2-6 kb. A total of 465 such autosomal exons were identified in the human genome (*i.e*., those that were relatively long and were less likely to have interference from other functional elements nearby) and used for further analysis (provided as a supplemental file; see Methods for further details).

The sensitivity of assuming the DFE inferred by Keightley and Eyre-Walker was evaluated by investigating how the reduction and recovery of nucleotide diversity due to BGS was affected by two very different DFE shapes – a DFE skewed strongly towards mildly deleterious mutations and another towards strongly deleterious mutations (Table S1). The primary determinant of BGS effects around functional elements was driven by the cross-over rate, for which we accounted. The DFE skewed towards strongly deleterious mutations predicted larger number of bases required for recovery (as expected from previous theoretical results). Note that while extremely strongly deleterious mutations have long range BGS effects, they reduce diversity only by a factor of ∼0.999 (in the human population), and do not segregate at high frequencies, and thus are unlikely to contribute to interference effects. Moreover, as all calculations were performed assuming a conservative absence of gene conversion, it is unlikely that there exist unaccounted for interference effects from other nearby exons.

**Table 1:**
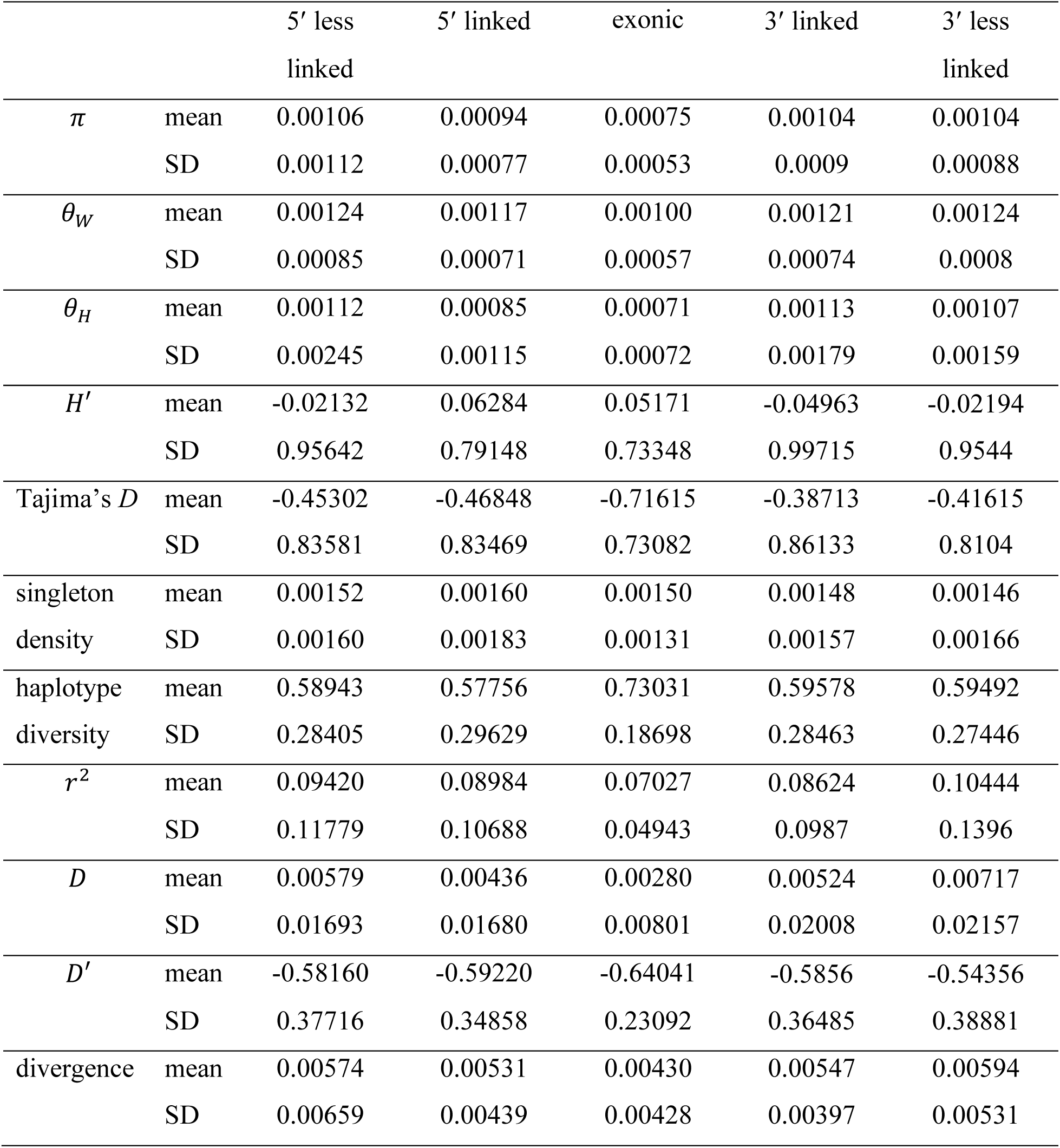
Means and variances of statistics from the empirical data for all 465 exons and their linked non-coding regions.

### The ABC approach

An ABC approach was employed to perform joint inference of parameters of demography and purifying selection while accounting for BGS effects. As BGS tends to distort genealogies such that inferences of recent population history could be biased, for the purpose of this study, recent size changes were specifically modeled and focused upon. More specifically, a simple single size change was modeled ∼200 generations ago (allowing for uncertainty in the age), which represents the Bantu population expansion (Schiffels and Durbin 2014), allowing estimation of the ancestral population size (*N_anc_*), current population size (*N_cur_*), and the precise time to change (*t* Figure 1a). Purifying selection was modeled using a DFE comprising four non-overlapping uniform distributions (Figure 1a), such that an *f*_0_proportion of all new mutations was neutral (2*N_e_s* = 0), an *f*_1_proportion was weakly deleterious (1 < 2*N_e_s* ≤ 10), an *f*_2_ proportion was moderately deleterious (10 < 2*N_e_s* ≤ 100), and an *f*_3_ proportion was strongly deleterious (2*N_e_s* ≥ 100). Note that *s* > 0 represents the selection coefficient against homozygotes and the effective population size (*N_e_*) here corresponds to the ancestral size. By sampling different combinations of *f*_i_ (such that *f*_*i*_ ∈ [0,1] ∀ *A* and ∑^*i*^ *f*_*i*_ = 1), all possible shapes of the DFE could be sampled (including bimodal DFEs). Thus, the inferred parameters of the DFE were the four proportions (*f*_*i*_) of new mutations in each selective class.

As ABC is a simulation-based method, all 465 exons were simulated using the forward time simulator SLiM (Haller and Messer 2019), with the specific lengths of exonic and intergenic/intronic regions, as well as their respective recombination and gene conversion rates (see Methods). Note that although the inference approach applied here is conceptually similar to that employed by Johri et al. (2020) to *Drosophila* populations, the simulations performed for the purpose of this work were tailored to the exons in the human genome and thus were newly performed. In addition, we have here newly added a consideration of gene conversion to the model. For each exon, statistics were calculated for three separate windows: (1) “functional” (comprising all sites in the exon), (2) “linked” (comprising *Π*_50_consecutive bases in the intergenic/ intronic region), and (3) “less linked” (comprising the subsequent set of *Π*_50_bases in the intergenic/intronic region; Figure 1b). A large number of statistics summarizing the means and variances of the site frequency spectrum, linkage disequilibrium (LD), and divergence for each window were employed when performing inference procedures (see Methods). The accuracy of inference was assessed by performing a leave-one-out cross-validation, whereby a single parameter combination was excluded from the priors while performing inference. All seven parameters (*N_anc_*, *N_cur_*, and *f*_0_–*f*_3_) were estimated sufficiently well (Figure 1c, Table S2), with the smallest errors in the proportion of neutral mutations (*f*_0_) and the ancestral population size, and highest errors in the estimation of the proportion of moderately deleterious mutations (*f*_2_) and the current population size.

Genomic analyses of sequencing data – such as those collected for the Yoruba population as part of the 1000 Genomes project (Auton et al. 2015) – are inevitably restricted to sites in the genome that are accessible to next-generation sequencing. In addition, summary statistics are frequently reported for regions outside of functional elements to avoid the effects of selection. Such filtering could potentially bias the values of statistics, particularly those associated with the variance of the statistics across exons. Indeed, when a filtering scheme replicating that of the 1000 Genomes project was employed on simulated data, a drastic increase in the variance of SFS-based statistics was observed post-filtering (Figures S2 and S3; Table S3), while almost no change was observed for statistics based on LD. Therefore, the sites excluded from the empirical data (*i.e.*, inaccessible sites and those belonging to functionally important regions smaller than 500 bp) were also excluded from the simulated data, decreasing the accuracy of our inference method almost by half (Table S2). Importantly, by mimicking the filtering of the empirical data in such an exact manner in the simulated data, the statistics observed in the YRI population across the 465 exons (as shown in Table 1) were well explained by the set of simulations employed for inference (Figures S4-S10).

### Inference and comparison to previous studies

Upon performing inference on the Yoruba population, a 6-fold population size increase was estimated that began ∼300 generations ago (with an ancestral and current size of 7,509 and 44,632 individuals, respectively). These estimates correspond well to previous neutral estimates of the recent history of the Yoruba population (Figure 2a). Interestingly, by jointly estimating population history and the DFE, our estimated shape of the deleterious DFE was highly skewed towards mild selective effects – ∼50% of all new mutations in the exonic regions investigated were estimated to belong to the effectively neutral class, ∼20% to the mildly deleterious class, and ∼30% to the moderately deleterious class. In addition, when dividing the 465 exons into two equal sets with high and low exonic divergence from the human ancestor, a much larger proportion of effectively neutral mutations was observed in the high-divergence set as expected (Figure 2b). These observations suggest that it is possible to infer the DFE with reasonable accuracy, and that the shape will depend upon the set of chosen exons (also see Campos et al. 2017).

**Figure 2:**
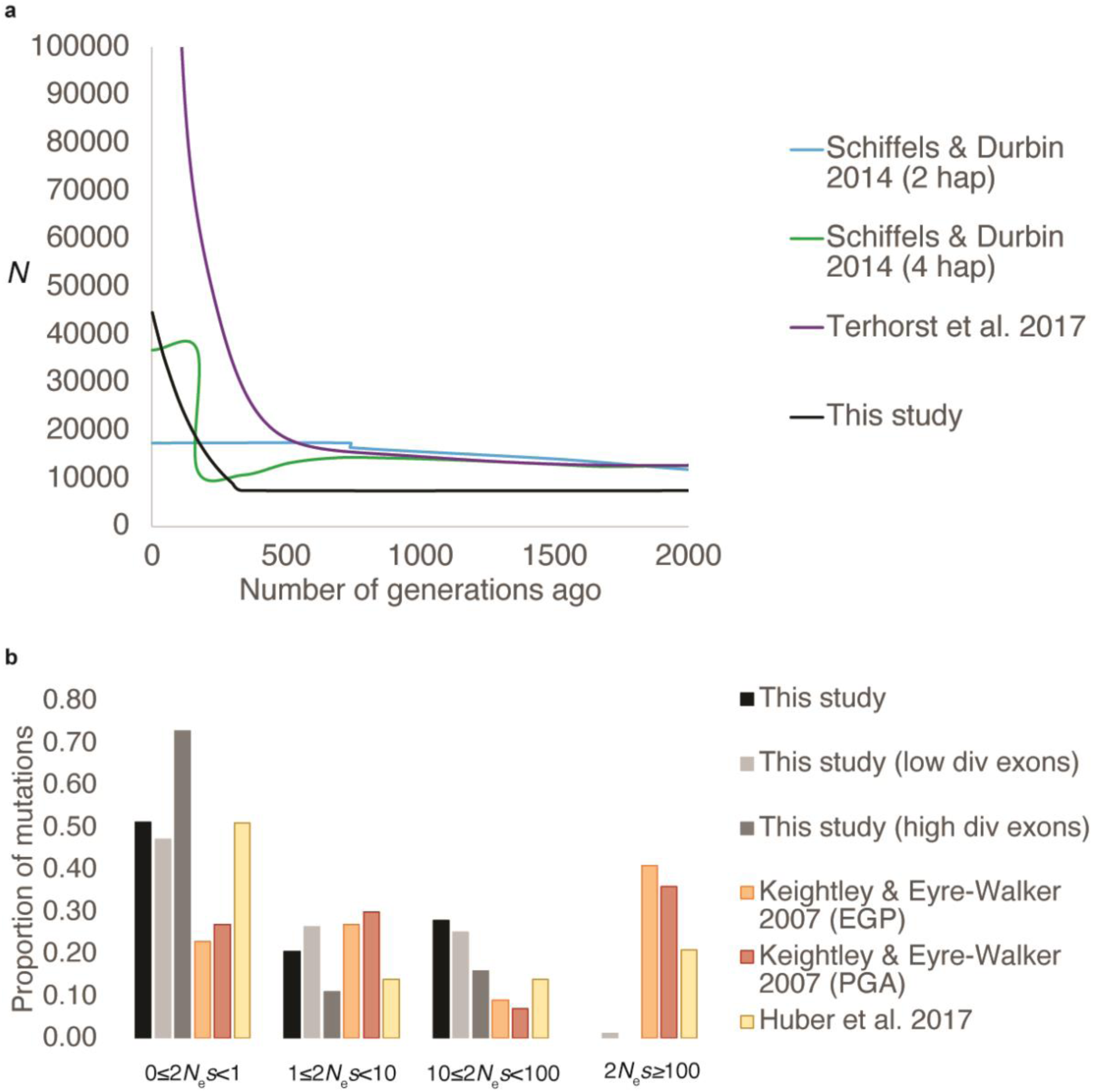
Inference of (a) recent population history and (b) the DFE of deleterious mutations in the Yoruba population. Inferences from the current study (using the 5′ intergenic/intronic regions) are shown in grey/black while those from previous studies are shown in the colored bars/lines. Note that the current population size predicted by Terhorst et al. (2017) is 356,990 and is not visible due to truncation of the *y*-axis. Also note that 2*N_e_s* for the purpose of the current study corresponds to 2*N_anc_^s^* as the scaling was performed with respect to the ancestral population size. 2 hap: refers to inference performed using a single diploid individual; 4 hap: refers to inference performed using 2 diploid individuals; EGP: Environmental Genome Project (https://egp.gs.washington.edu/); PGA: Programs for Genomic Applications (https://pga.gs.washington.edu/).

This approach did not identify an appreciable proportion of strongly deleterious mutations amongst these selected exons, though there is of course some uncertainty around these estimates (presented as the posterior distributions provided in Table S4 and Figure S11). Notably however, as previous studies have generally assumed a gamma distribution of the deleterious DFE, it is also possible that constraints of the gamma distribution have resulted in the estimation of more mutations in the strongly deleterious class. Moreover, the DFE estimated by Keightley and Eyre-Walker (2007; 2009) was estimated based on a set of genes selected either because loss-of-function mutations in those genes cause severe diseases (EGP data), or because those genes underly inflammatory responses (PGA dataset) in humans. Such genes are more likely to be highly conserved and thus to have more strongly deleterious mutations. Huber et al. (2017) used a wider set of genes and obtained a DFE more skewed towards effectively neutral mutations, with a very similar shape as that obtained by the present study (Figure 2b). Further, these observed differences in the DFE could also reflect differences in the DFE of different populations – the present study was conducted on the Yoruba population (the same dataset analyzed by Huber et al.), while the DFE in Keightley and Eyre-Walker (2007) was calculated from an African-American population. Finally, as our method could only be applied to exons located in sparser regions of the human genome, limited to 465 in number, it is possible that the difference in the estimated DFE from Huber et al. (2017) is due to the difference in selective constraints experienced by the selected group of exons vs. all exons.

### Model violations and fit

In order to find a sufficient number of exons that were appropriately distant from other functional elements, we excluded exons that were near phastCons elements of lengths larger than 500 bp (see Methods for details). Thus, hitchhiking effects (due to selective sweeps and/or BGS) generated by smaller phastCons elements were not accounted for in the priors. Note that most phastCons elements are extremely small in length, with 50%, 90%, and 99% of all phastCons elements being less than 10 bp, 32 bp, and 132 bp respectively (Table S5). Theoretical calculations using Equations 1-5 and assuming a DFE skewed towards mildly deleterious mutations (*f*_0_ = 0.1; *f*_1_ = 0.7; *f*_2_ = 0.1; *f*_3_ = 0.1) demonstrates that BGS effects generated by such small functional elements are extremely minor (with *B*=0.993-1.0; Table S6) and are thus highly unlikely to cause unaccounted for interference effects.

Another potential caveat of our analysis is the assumption that ancestral alleles have been accurately inferred by previous studies. Keightley and Jackson (2018) noted two consequences of ancestral allele misidentification on biasing estimation of summary statistics. Firstly, when parsimony methods are used to infer the derived allele, filtering of sites can lead to a decrease in levels of diversity. As the 1000 Genomes data used multiple outgroups to polarize SNPs, it will likely result in stringent filtering criteria (possibly removing sites that have a high mutation rate). Hence, while such a bias may lead to under-estimation of population sizes, a comparison of our estimates to those from previous studies is justified (as other studies are also using the same ancestral alleles to polarize SNPs). The second issue noted by the authors is that parsimony methods can result in an over-estimation of high frequency derived alleles. However, they observed that the unfolded SFS from the 1000 Genomes dataset is very similar to what they obtained using their maximum likelihood approach (corrected for the mis-identification), unless they restricted it to CpG sites. As we are not particularly looking at CpG sites alone (and as noted above, we are likely throwing a number of those out), our SFS should not be biased. In order to formally test this, the following analysis was performed to evaluate the sensitivity of our results to mis-specification of the ancestral state. As CpG sites comprise of less than 1% of the human genome (Babenko et al. 2017), it was assumed that ∼1% of all derived singletons were falsely polarized, and thus were randomly re-assigned to an allele frequency of 99%. Note that as not all CpG sites will have segregating derived singletons, this example assumes that many more sites have a mis-specified ancestral state than is likely to occur in real data. The accuracy of inference of parameters related to the demographic history were almost entirely unaffected by this mis-specification (Table S7). However, there was an under-estimation of the fraction of mutations in the weakly deleterious class and a corresponding over-estimation of the moderately deleterious class (Table S7).

Finally, a potential caveat concerning the inferences performed in this study is the assumption of a common mutation rate across the simulated regions. Region-specific mutation rates estimated from the identification of *de novo* mutations in humans were therefore simulated to assess the magnitude of generated bias in our inference method. Again, while inference of parameters of the demographic history were unaffected, there was a slight under-estimation of the fraction of mildly deleterious mutations when mutation rate heterogeneity was neglected (Table S7). Thus, although a large class of mildly deleterious mutations was inferred from the human data, the proportion of weakly deleterious mutations may be even higher. Despite this caveat, our inferred model fits the data exceptionally well for all classes (functional, linked, less-linked) and across all 465 exons (Figures 3 and S12-S14). This fit was evaluated by simulating the best-fit model 10 times, and comparing the distribution of all the summary statistics with those obtained from the empirical data. As can be seen in Figure 3, predicted patterns of LD, the SFS, and divergence match the empirical data well. It should be noted that the figure compares the entire distribution of statistics for all 465 exons between the best-fitting model and the empirical data – a comparison that is usually restricted to mean values of summary statistics, and thus suggests an overall excellent fit between the estimated best model and real data. Importantly, despite the strong fit of our inferred model to the data, it is very likely that additional parameter combinations under alternative models (including a more complex demographic history) could also be fit to the data (as discussed in Johri et al. 2022a). As such, this model (as with any model fitting exercise) should only be viewed as a viable model rather than, of course, as a ’correct’ model.

**Figure 3:**
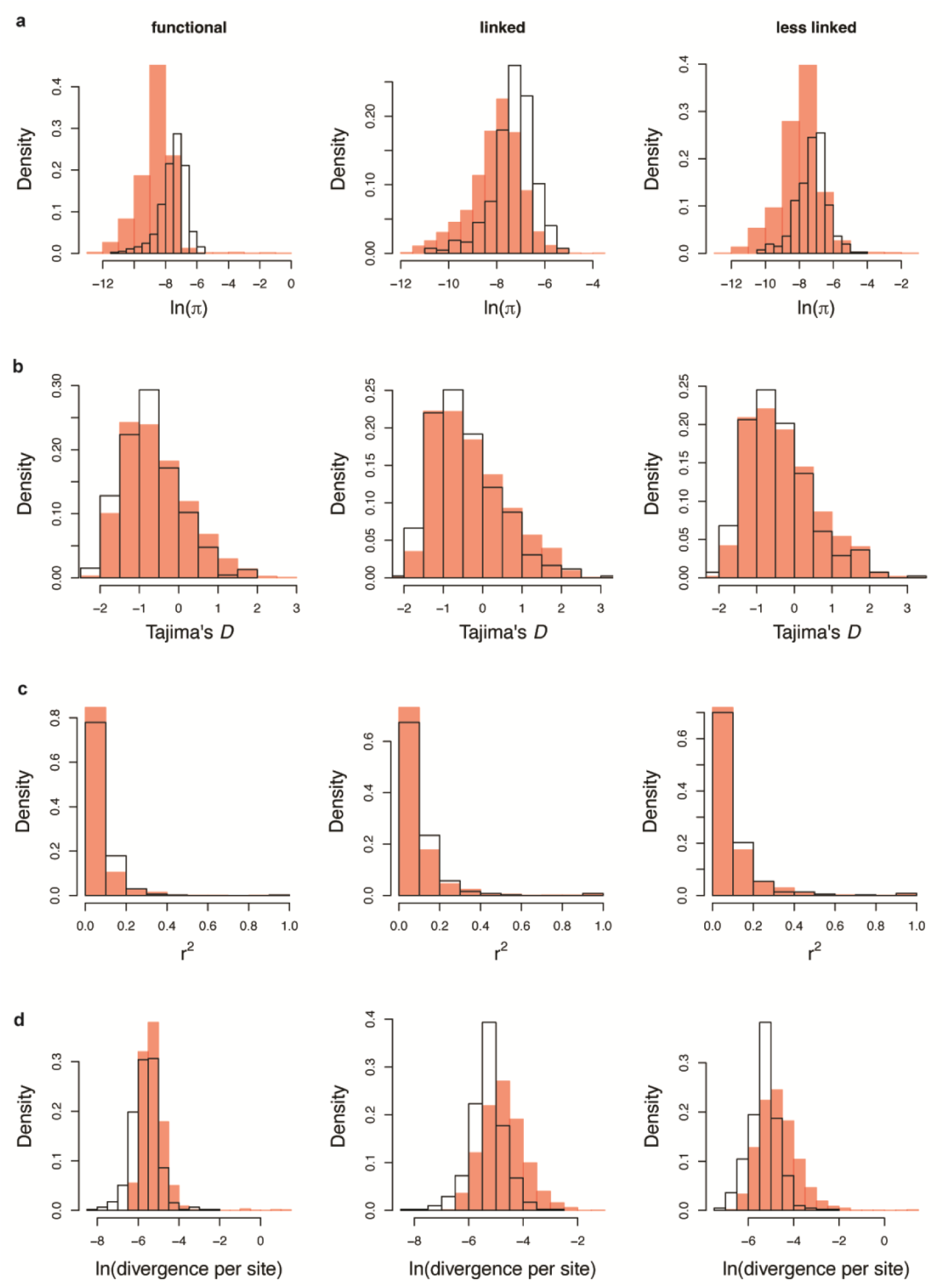
Fit of the best model inferred by our method to the empirical data, as shown by the distribution of (a) nucleotide diversity, (b) Tajima’s *D*, (c) *r*^2^, and (d) divergence per site, across the 465 exons, for each of the three windows: functional, linked and less linked intergenic/intronic regions. The simulated best model (with 10 replicates) is shown in red, while the observed empirical distributions of the same statistics in the YRI population are shown in the white distributions.

### Evaluating the identifiability of a beneficial mutational class

As beneficial mutations are expected to be rare and only episodically reach fixation, they were not part of the baseline model fit to the data, which was instead focused upon commonly and continuously acting evolutionary processes. Nonetheless, the identification of beneficial mutations is of great interest, and thus the effects of a model violation consisting of various rates and strengths of recurrent positive selection within the context of the fit baseline model was evaluated. The proportion of new beneficial mutations (*f*_*pos*_) was assumed to be 0.1%, 1%, or 5%, and the distribution of fitness effects of beneficial mutations was modelled to be exponentially distributed with mean *s*_*b*_, such that 2*N_e_s*_*b*_ = 10, 100, or 1000, where *s*_*b*_ > 0 is the selection coefficient representing selection favoring homozygotes, and all mutations were assumed to be semidominant. Combinations of the above parameters yielded nine different evolutionary scenarios ranging from weak and infrequent to common and strong positive selection.

Interestingly, when 0.1% or 1% of new mutations are beneficial and the strength of positive selection is weak or moderate (2*N_e_s*_*b*_ = 10 or 100), there is almost no difference between the distribution of statistics across the 465 exons in the absence vs. presence of positive selection (Figures S15-S23). This observation is consistent with results from *Drosophila melanogaster* (Johri et al. 2020), and suggests a general inability to identify this class of mutations, if present. However, when positive selection is common (*f*_*pos*_ = 1 − 5%) and strong (2*N_e_s*_*b*_ = 1000), the distribution of statistics including Tajima’s *D*, *r*^2^, and divergence do not resemble observed empirical distributions (Figures 4 and S15-S23). Therefore, while strong and frequent positive selection is inconsistent with empirical data, weak/moderate infrequent positive selection remains consistent with observed patterns of variation, though this addition does not improve the fit. This observation emphasizes the peril of naively fitting models of positive selection to data while neglecting common evolutionary processes (see Johri et al. 2022c), as well as the difficulty in being able to accurately infer the proportion and DFE of new beneficial mutations. More generally however, it will be of interest in the future to evaluate whether a joint inference approach that explicitly includes a class of beneficial mutations can be successful.

**Figure 4:**
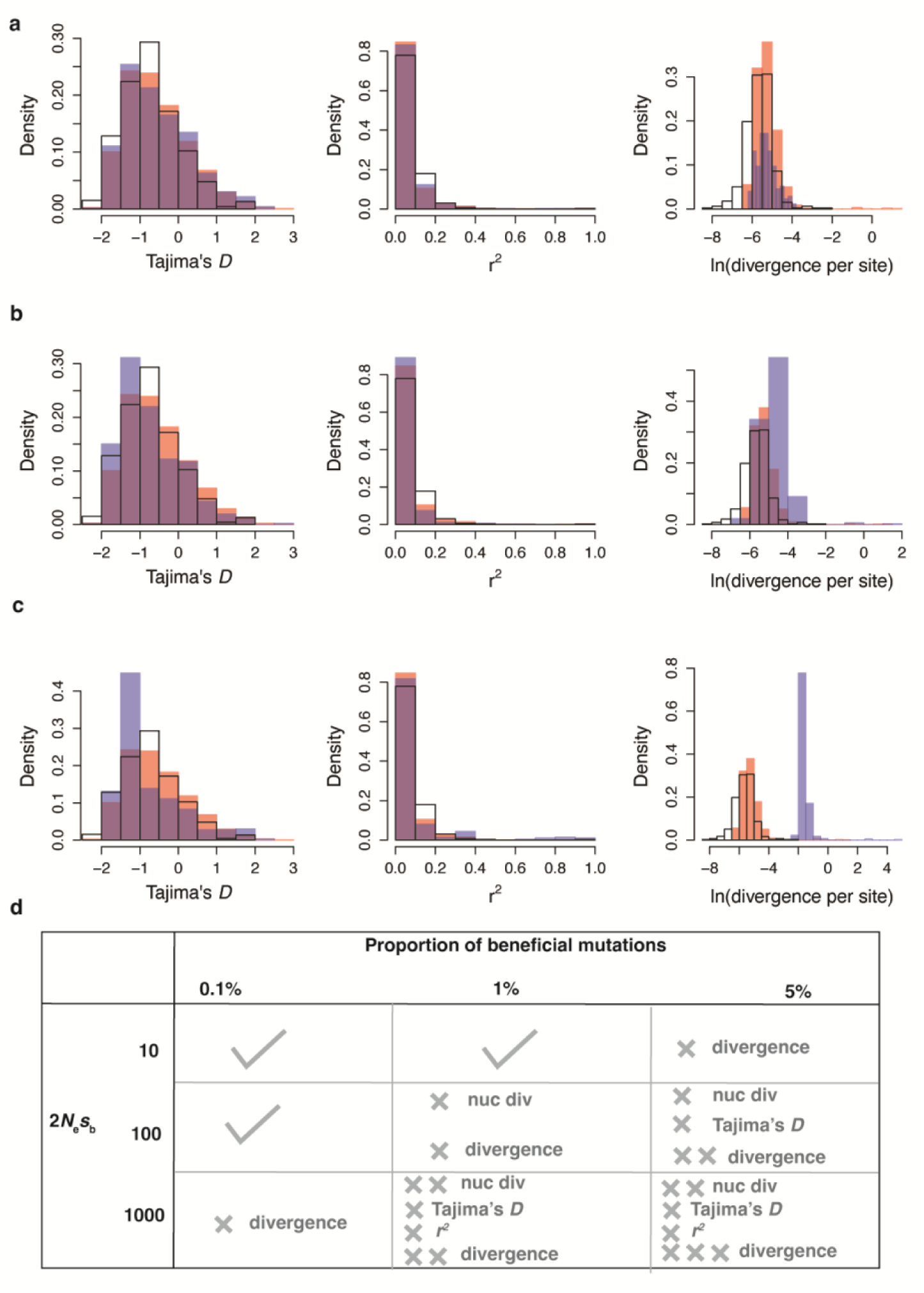
Fit of the estimated best model to the empirical data in the presence of positive selection. (a-c) Distribution of Tajima’s *D*, *r*^2^, and divergence per site across the 465 exons (only in the “functional” windows) for the best-fitting model (in red), the best-fitting model with positive selection (in blue), and their overlap (in purple). The distribution of the empirical data is shown in the white distributions. Examples of varying extents of positive selection are shown: (a) infrequent (*f*_*pos*_ = 0.1%) and weak (2*N_e_s* = 10), (b) moderately frequent (*f*_*pos*_ = 1%) and moderately strong (2*N_e_s* = 100), and (c) common (*f*_*pos*_ = 5%) and strong (2*N_e_s* = 1000). (d) A grid depicting the fit of varying extents of positive selection to the data with a check mark indicating that the addition of positive selection does not worsen the fit of the model to the data, and the number of ’×’ marks indicating the severity of the mis-fit to the calculated statistics generated by the addition of positive selection.

### Closing Thoughts

Despite some important methodological differences amongst approaches, one commonality that has emerged in the study of the Yoruba population is the presence of an appreciable class of weakly deleterious (1 < 2*N_e_s* ≤ 10) mutations. This observation has a few noteworthy implications. Firstly, the BGS effects arising from mildly deleterious mutations cannot be accounted for by a simple rescaling of effective population size, as these mutations will result in a significant skew towards rare alleles; this may in turn strongly bias demographic inference when unaccounted for (Ewing and Jensen 2016; Johri et al. 2021). Secondly, weakly deleterious mutations in regions of low recombination can result in associative overdominance, which could lead to an increase in both nucleotide diversity and LD (Zhao and Charlesworth 2016; Becher et al. 2020; Gilbert et al. 2020). Additionally, the linked effects of very weakly deleterious mutations (1 < 2*N_e_s* < 2.5) are still not well understood (though see Charlesworth 2022), and thus their common presence in such inference suggests the need for further study of these weak selection effects.

Finally, as the human genome is characterized by a small fraction (< 10%) of functional sites (Siepel et al. 2005), and indeed is amongst the best-annotated and best-studied genomes to date, this species probably represents a case for which the joint inference of demography with the DFE is least critical. In other words, in functionally dense genomes in which neutral sites free of BGS effects may be difficult to identify or may not exist at all, as well as in less well-studied species in which functional elements may not be fully annotated for the purposes of exclusion when performing demographic inference, this type of joint inference will be critical for accurate estimation. This disparity is partly evidenced by comparing joint inference performed in *D. melanogaster* and in humans. In the former, the incorporation of BGS effects into the joint inference scheme led to considerably lower estimates of population growth and higher proportions of weakly deleterious mutations relative to studies utilizing two-step inference approaches (Johri et al. 2020), whereas in humans the joint inference estimates provided here are relatively similar to previous two-step estimates.

That said, earlier studies in humans as well as model organisms such as *D. melanogaster* (e.g., Beichman et al. 2017; Garud et al. 2021) have shown that the specific models of demography that have been fit previously to these populations do not recapitulate all aspects of the data. Specifically, when the inferred models are simulated, they explain certain aspects of the data, but poorly fit others (*e.g.,* linkage disequilibrium). Conversely, we have here shown that incorporating the specific details of genome architecture with locus-specific recombination rates, employing a statistical approach that can account for multiple aspects of the data, and jointly inferring population history with the DFE utilizing BGS expectations, results in a remarkably good fit to all aspects of levels and patterns of variation and divergence. This once again highlights the importance of constructing an appropriate evolutionary baseline model for genomic analysis, and of relaxing common but poorly supported inference assumptions.

## METHODS

### Data

This study was based on the human reference genome hg19/GRCh37 and its corresponding resources. In brief, the human reference genome (hg19) was downloaded from the UCSC Genome Browser (accession number: GCA_000001405.1; Church et al. 2011); a catalogue of common genetic variation in the Yoruba population was obtained from the 1000 Genomes Phase 3 (Auton et al. 2015) together with information about genome accessibility to next-generation sequencing (as determined by the “tgpPhase3AccessibilityStrictCriteria” track of the UCSC Table Browser); ancestral alleles as determined by the six-way primate EPO alignments were downloaded from Ensembl (release 74; Flicek et al. 2014; Cunningham et al. 2022); gene annotations (including exon start and end positions) were downloaded from the NCBI Human Genome Resources archive (Sayers et al. 2022); annotations for small nucleolar and micro RNAs (“sno/miRNAs”) as well as conserved elements identified based on the 100-way PhastCons score (“phastConsElements100way”; Siepel et al. 2005; Pollard et al. 2010) were downloaded from the UCSC Table Browser; and population-specific recombination rates were obtained from the HapMap project (“hapMapRelease24YRIRecombMap”; Altshuler et al. 2005). The URLs for file downloads are provided in Table S8.

### Selecting a set of human exons for analysis

For every exon in the human genome, we calculated the decay of nucleotide diversity at linked neutral sites caused by BGS, taking into account the specific exon length and recombination rate (assuming the rate of gene conversion to be zero in order to be conservative). This was done analytically using equations 3a and 3b derived in Johri et al. (2020), and presented as equations 1-5 in the Results section. The DFE was assumed to follow that inferred by Keightley and Eyre-Walker (2007): *f*0=0.22, *f*1=0.27, *f*2=0.13, and *f*3=0.38, representing the proportion of new mutations belonging to the neutral, weakly deleterious, moderately deleterious, and strongly deleterious classes, respectively. Nucleotide diversity was predicted at sites 1 to 100,000 bp away from the end of each exon and a logarithmic function was fit such that *Π* = *slope* × ln(*x*) + *intercept*, where *x* is the distance of the site from the functional region in base pairs. The values of *slope* and *intercept* were used to calculate the expected number of base pairs required for a 50% recovery of *Π* (referred to as *Π*_50_). The script to perform such analytical calculations can be accessed at: https://github.com/paruljohri/Joint_Inference_DFE_demog_humans/blob/main/selecting_exons/a dd_numbp50_to_exons.py. The distance between every exon and its nearest functional element (*i.e*., all neighboring exons, as well as sno/miRNAs and phastCons elements larger than 500 bp) was calculated, and exons with a distance greater than 4 × *Π*_50_ kept for further analysis. In addition, only exons that were 2-6 kb in length were selected (in order to observe significant BGS effects). This procedure yielded a total of 465 exons with recombination rates within 0.5-10cM/Mb. Note that the selected exons were not restricted to single-exon genes.

### Modeling the simulation framework for ABC

Each of the 465 exons was simulated using SLiM v.3.1 (Haller and Messer 2019), and was comprised of a functional region of the length of the exon, with a single linked intergenic/intronic region of size 4 × *Π*_50_. Intergenic/intronic regions were assumed to be neutral, whereas exonic regions experienced purifying selection given by a discrete DFE comprised of four non-overlapping uniform distributions, with *f*_0_, *f*_1_, *f*_2_and *f*_3_representing the proportion of new mutations belonging to the neutral, weakly deleterious, moderately deleterious, and strongly deleterious classes, respectively. Simulations were performed using a constant mutation rate of 1.25 × 10^−8^ per site per generation (Kong et al. 2012) and region-specific crossing over rates obtained from the HapMap project (Altshuler et al. 2005) as indicated in the Data section and Table S8, utilizing the average crossing over rate across the exonic and corresponding intergenic/intronic regions (both 5′ and 3′ intergenic/intronic) for each exon.

#### Modeling gene conversion

The rate of non-crossover gene conversion has been estimated to be 5.9 × 10^−6^ per site per generation (Palamara et al. 2015; Williams et al. 2015), with tract lengths found to be between 55-290 bp (Jeffreys and May 2004) and between 100-1000 bp (Williams et al. 2015). In humans and mice, crossover recombination events (COs) are ∼5–15 times less frequent than non-crossovers (NCOs), but their conversion tracts are ∼2–8 times longer (Jeffreys and May 2004) and most of these events occur in recombination hotspots (McVean et al. 2004). Although the mean rate of gene conversion per site (*i.e.*, the probability that any given site is affected by the process of gene conversion) can be estimated with confidence and is consistent across the studies mentioned above, it is quite difficult to disentangle the tract length from the rate of initiation of gene conversion. Moreover, previous studies have found that gene conversion rates are correlated with the rate of crossing over in humans (Padhukasahasram and Rannala 2013; Glémin et al. 2015; Palamara et al. 2015). We thus assume that tract lengths are geometrically distributed (as modeled in SLiM) with a mean of 125 bp (Jeffreys and May 2004), and that gene conversion rates are 5 times those of recombination rates, while maintaining the average rate of gene conversion of 5.9 × 10^−6^per site per generation.

#### Demographic history

To correct for confounding effects of BGS on population history, a simple demographic history comprised of a single, recent population size change was modeled. As Gutenkunst et al. (2009) and Gravel et al. (2011) fit a single size change model that yielded a size change relatively long ago (∼6-8k generations), those models were not used to parametrize the model in this study. Instead, we based our model on previous studies that have estimated a recent increase in population size of the Yoruba population, corresponding to the Bantu expansion, with the estimated expansion occurring ∼200 generations ago. Specifically, Tennessen et al. (2012) estimated the time of change to be 205 generations ago (corresponding to 5,115 years ago with a generation time of 25 years), Schiffels and Durbin (2014) estimated the time of change to be 200 generations ago (corresponding to 6,000 years ago assuming a generation time of 30 years), and Terhorst et al. (2017) estimated that the growth in the population began 1,724 generations ago and significantly increased around 517 generations ago (corresponding to 50k and 15k years ago assuming a generation time of 29 years). The YRI population was thus simulated to be under equilibrium until a size change (exponential increase or decrease) occurred ∼200 generations ago (referred to as *τ*) with uncertainty modeled around this age (Figure S24). The ancestral (*N_anc_*) and current (*N_cur_*) population sizes were inferred using ABC (see below).

### ABC

A total of seven parameters were inferred using ABC: *f*_0_, *f*_1_, *f*_2_, *f*_3_, *N_anc_*, *N_cur_*, and *t*.

The *f*_*i*_ were randomly sampled in increments of 0.05 between 0 and 1, *i.e*., *f*_*i*_ ∈ {0.0, 0.05, 0.1, …, 0.95, 1.0} such that 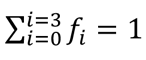1. Both *N_anc_* and *N_cur_* were sampled from log uniform distributions between 5,000 and 50,000 diploid individuals. A total of 2000 parameter combinations were simulated. Simulations for each parameter combination were rescaled to a different extent, determined as follows. In order to avoid simulating extremely small population sizes and having a very large rescaling factor, rescaling was restricted to a maximum of 200-fold and a minimum of 5,000 individuals, *i.e*., 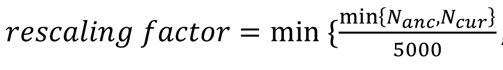. For each parameter combination, the 465 exons with their specific lengths, intergenic/intronic region, cross-over and non-crossover rates were simulated for a burn-in period of 10*N_anc_* generations plus an additional 4*N_e_*_*anc*_ generations (in order to estimate the rate of divergence post burn-in) after which there was an exponential size change for *τ* generations. Fifty diploid individuals were sampled at the end of each simulation.

#### Calculation of statistics from simulated data

For each exon, three non-overlapping windows were defined: (1) “functional” (comprised of all sites in the exonic region), (2) “linked” (sites within [0, *Π*_50_] bases linked to the exon, with 5′ and 3′ being designated separately), and (3) “less linked” (sites within (*Π*_50_, 2*Π*_50_] bases linked to the exon, with 5′ and 3′ being designated separately). Next, any sites deemed inaccessible in the 1000 Genomes Phase 3 data were excluded (see Data section above) and sites in the intergenic/intronic regions that were annotated to be functionally important (*i.e.*, phastCons elements) that were smaller than 500 bp were also excluded. Pylibseq v.0.2.3 (Thornton 2003) was used to obtain the means and standard deviations of the following statistics from both the unfiltered and filtered simulated data: nucleotide diversity (*Π*), Watterson’s *θ* (*θ*_*W*_), statistics that capture the relative proportion of high and intermediate frequency alleles (*θ*_*H*_,), statistics that capture the relative proportion of rare alleles of the SFS (Tajima’s *D*, singleton density), and statistics that summarize the LD patterns (haplotype diversity, *r*^2^, *D*, *D*′). Together with divergence (see below) these amounted to a total of 66 summary statistics that were employed to perform inference using the ABC method. It should be noted that while ABC-based approaches can suffer from the “curse of dimensionality”, *i.e*., ABC inference can become inaccurate and unstable if an extremely large number of statistics are employed (Beaumont 2010), excluding statistics always resulted in a reduction of accuracy in our study. Moreover, different statistics from different windows were important to accurately predict different parameters (see Johri et al. 2020 for a detailed analysis). Therefore, all 66 summary statistics were used for inference.

Divergence was calculated using the number of substitutions (as provided by SLiM) that occurred after the burn-in period of 10*N_anc_* generations. As a rate of substitutions was obtained from the simulations, these rates were converted to divergence values as follows. The total number of fixed substitutions per site (*div_rate_*) was calculated from the simulations over the course of 4*N_anc(scaled)_* + 200 generations for each parameter combination. Divergence values (*div*) for each parameter combination were then obtained using

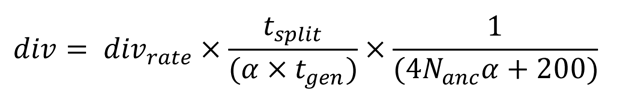

where *t*_*split*_ is the time since the split between chimpanzees and humans, which was assumed to be a minimum of 6 (Nachman and Crowell 2000) and a maximum of 12 million years ago (Chintalapati and Moorjani 2020) and the mean of these values was used when performing final inference. The generation time (*t_hen_*) in humans was assumed to be 25 years following Gutenkunst et al. (2009) and *α* is a scaling factor (which was different for each parameter combination). Thereby, the sites that were excluded when calculating statistics from single nucleotide polymorphisms (SNPs) were also excluded when calculating divergence from simulated data. Note that as divergence per site was calculated by multiplying the rate of fixation of mutations in simulated data with the total number of generations to the ancestor, and as filtering of sites resulted in some regions having very few accessible sites (*e.g*., 4-7), by chance some replicates had higher values, which resulted in values of divergence per site > 1 in this extreme parameter space (as can be seen in Figures 3 and 4).

#### Calculation of statistics from empirical data

Summary statistics were calculated from 50 YRI individuals (25 males and 25 females) selected at random from the 1000 Genomes Phase 3 data (Auton et al. 2015; and see Table S9 for the list of individuals), using only sites located in strictly accessible regions and outside of phastCons elements (see Data section above). Similar to the simulated data, pylibseq v.0.2.3 (Thornton 2003) was used to calculate the means and standard deviations of the 66 summary statistics (as outlined above), based on the 81% of sites retained after filtering (89% in exons, 80% in 5′ intergenic/intronic regions, and 79.4% in 3′ intergenic/intronic regions; Figure S25). Thereby, divergence was calculated based on fixed differences between reference and ancestral alleles that were non-polymorphic in the YRI dataset. Final values of all statistics obtained from the 50 randomly selected diploid YRI individuals were very similar to those obtained using all individuals (Table S10).

### ABC inference

The ABC approach was executed using the R package “abc” (Csilléry et al. 2012). A correction for the non-linear relationship between the parameters and the statistics was employed using the “neural net” regression method with the default parameters provided by the package. A 100-fold leave-one-out cross-validation was performed in order to determine the performance and accuracy of inference for the following values of tolerance: 0.05, 0.08 and 0.1. As inference was most accurate with a tolerance of 0.08 (Table S2), this value was employed for inference of final parameter values, *i.e.*, 8% of all simulations were accepted by ABC to estimate the posterior probability of parameter estimates. Final point estimates for each parameter were calculated by performing inference 50 times and taking the mean of the (50) weighted medians of the posterior estimates.

### Simulations with mutation rate heterogeneity

Sex-averaged mutation rate maps for humans were obtained from Francioli et al. (2015) (https://www.nlgenome.nl/menu/main/app-download; last accessed Sep 22, 2022). As rates were provided for each nucleotide (*i.e*., A to C, A to G, etc.), the nucleotide composition of each exon was determined separately for the 5’ intergenic/intronic, exonic, and 3’ intergenic/intronic regions to obtain region-specific mutation rates. Specifically, for each exon, the average rate of the three regions multiplied by a mean mutation rate of 1.25 × 10^−8^per site per generation (as rates were normalized with respect to this mean mutation rate) was used for simulations.

### Simulations of the best-fitting model

When simulating the best-fitting model, the best estimates (weighted median) of each parameter were used. Ten independent replicates of each of the 465 exons were simulated and the distribution of all statistics (post-filtering) for each window was compared to the corresponding empirical data.

### Simulations with positive selection

When simulating the best-fitting model with positive selection, the best estimates (weighted median) of each parameter were used. To test the effect of recurrent selective sweeps, beneficial mutations were assumed to be a fraction *f*_*pos*_ of all new mutations in exonic regions and their fitness effects were assumed to follow an exponential distribution with mean *s*, such that 2*N_ancs_* = 10, 100, or 1000. The fitness effects of the remaining exonic mutations (1 − *f*_*pos*_) followed the estimated DFE (comprising neutral and deleterious mutations).

## ACKNOWLEDGMENTS

We thank Brian Charlesworth for providing helpful comments and suggestions on the manuscript, and John Terhorst for providing us with the exact values of population history obtained in Terhorst et al. 2017. This research was conducted using resources provided by Research Computing at Arizona State University (http://www.researchcomputing.asu.edu) and the Open Science Grid, which is supported by the National Science Foundation and the U.S. Department of Energy’s Office of Science. This work was funded by National Institutes of Health grant R35GM139383 to JDJ.

## DATA AVAILABILITY

All scripts used to perform the research in this study have been made available at https://github.com/paruljohri/Joint_Inference_DFE_demog_humans. The final set of exons used in the study, along with their mutation and recombination rates, are provided as a supplemental file (single_exons_465_suppfile.xlsx).

## Notes

https://github.com/paruljohri/Joint_Inference_DFE_demog_humans

### Competing Interest Statement

The authors have declared no competing interest.

