## Supplemental Tables and Figures for "Developing an evolutionary baseline model for humans: jointly inferring purifying selection with population history"

**SUPPLEMENTAL FIGURES AND TABLES**

**Table S1**: The nucleotide diversity with BGS relative to that in the absence of selection (*B*), and the expected number of base pairs ($\pi_{50}$) required for a 50% recovery of diversity around exons of varying lengths, with varying rates of recombination and three different DFEs.

| **Exon length (bp)** | **Recombination rate**[1] | **DFE1**[2] | | **DFE2**[3] | | **DFE3**[4] | |
|---|---|---|---|---|---|---|---|
| | | $\pi_{50}$ | *B* | $\pi_{50}$ | *B* | $\pi_{50}$ | *B* |
| 2000 | 0.1 × mean | 48553 | 0.98 | 51201 | 0.98 | 79630 | 0.98 |
| | mean | 4000 | 0.98 | 4257 | 0.98 | 6420 | 0.99 |
| | 10 × mean | 38 | 0.99 | 48 | 0.99 | 197 | 0.99 |
| 4000 | 0.1 × mean | 58495 | 0.97 | 61926 | 0.97 | 98994 | 0.98 |
| | mean | 5012 | 0.97 | 5311 | 0.97 | 7774 | 0.98 |
| | 10 × mean | 114 | 0.98 | 138 | 0.98 | 432 | 0.98 |
| 6000 | 0.1 × mean | 70689 | 0.96 | 75104 | 0.96 | 123060 | 0.97 |
| | mean | 5903 | 0.96 | 6235 | 0.96 | 8946 | 0.97 |
| | 10 × mean | 199 | 0.98 | 233 | 0.98 | 650 | 0.98 |

[1] The mean recombination rate is $1 \times 10^{-8}$ per site per generation. [2] DFE1 refers to the Keightley and Eyre-Walker (2007) DFE with $f_0 = 0.22, f_1 = 0.27, f_2 = 0.13, f_3 = 0.38$. [3] DFE2 is skewed towards mildly deleterious mutations with $f_0 = 0.5, f_1 = 0.25, f_2 = 0.15, f_3 = 0.1$. [4] DFE3 is skewed towards strongly deleterious mutations with $f_0 = 0.1, f_1 = 0.15, f_2 = 0.25, f_3 = 0.5$.

**Table S2**: Prediction error of all parameters with and without filtering sites when performing inference using the ABC method. A 100-fold cross-validation was performed by excluding a single parameters combination at a time, using a tolerance of 0.05, 0.08 and 0.1, with a total of 2000 parameters combinations. Simulated sites were subject to the same filters as those applied to the empirical YRI dataset.

| | Tolerance | $f_0$ | $f_1$ | $f_2$ | $f_3$ | $N_{anc}$ | $N_{cur}$ | time |
|---|---|---|---|---|---|---|---|---|
| Without filtering | | | | | | | | |
| | 0.05 | 0.0337 | 0.1704 | 0.3499 | 0.1356 | 0.0088 | 0.2259 | 0.1286 |
| | 0.08 | 0.0354 | 0.1275 | 0.2817 | 0.1081 | 0.0113 | 0.1566 | 0.0995 |
| | 0.1 | 0.0321 | 0.1004 | 0.2206 | 0.0893 | 0.0115 | 0.1539 | 0.0757 |
| With filtering | | | | | | | | |
| 5′ | 0.05 | 0.0858 | 0.2086 | 0.2990 | 0.1479 | 0.0141 | 0.2926 | 0.1738 |
| | 0.08 | 0.0533 | 0.1857 | 0.3123 | 0.1559 | 0.0225 | 0.1853 | 0.1535 |
| | 0.1 | 0.0576 | 0.1644 | 0.2581 | 0.1094 | 0.0188 | 0.2014 | 0.1459 |
| 3′ | 0.05 | 0.0580 | 0.2567 | 0.4904 | 0.1836 | 0.0232 | 0.4278 | 0.3084 |
| | 0.08 | 0.0585 | 0.2036 | 0.3444 | 0.1364 | 0.0346 | 0.3276 | 0.2301 |
| | 0.1 | 0.0343 | 0.1925 | 0.3168 | 0.1195 | 0.0314 | 0.2957 | 0.1355 |

**Table S3**: Mean (across the first 1000 simulated parameter combinations) of the mean (across 465 exons) and variance (across the 465 exons) of all statistics with and without filtering sites. As shown, across different demographic histories and DFEs, there is an overall change in multiple statistics once filtering is employed.

| | | exonic | | $5'$ linked | | $5'$ less linked | |
|---|---|---|---|---|---|---|---|
| | | All sites | Filtered sites | All sites | Filtered sites | All sites | Filtered sites |
| $\pi$ | Mean | 0.00032 | 0.00100 | 0.00085 | 0.00113 | 0.00085 | 0.00176 |
| | SD | 0.00017 | 0.01094 | 0.00064 | 0.00118 | 0.00064 | 0.00770 |
| $\theta_W$ | Mean | 0.00042 | 0.00132 | 0.00088 | 0.00117 | 0.00089 | 0.00183 |
| | SD | 0.00016 | 0.01431 | 0.00049 | 0.00102 | 0.00049 | 0.00760 |
| $\theta_H$ | Mean | 0.00025 | 0.00077 | 0.00078 | 0.00104 | 0.00079 | 0.00162 |
| | SD | 0.00025 | 0.00880 | 0.00100 | 0.00162 | 0.00100 | 0.00824 |
| $H'$ | Mean | 0.19513 | 0.19467 | 0.08240 | 0.08270 | 0.08371 | 0.08374 |
| | SD | 0.55778 | 0.55884 | 0.78173 | 0.78128 | 0.77473 | 0.77453 |
| Tajima's $D$ | Mean | -0.59899 | -0.59746 | -0.13480 | -0.13464 | -0.13106 | -0.13096 |
| | SD | 0.72250 | 0.72294 | 0.84343 | 0.84360 | 0.84360 | 0.84365 |
| Singleton density | Mean | 0.00060 | 0.00187 | 0.00091 | 0.00121 | 0.00091 | 0.00188 |
| | SD | 0.00043 | 0.02074 | 0.00111 | 0.00179 | 0.00110 | 0.01001 |
| Haplotype diversity | Mean | 0.49253 | 0.49297 | 0.54031 | 0.54035 | 0.54074 | 0.54079 |
| | SD | 0.19867 | 0.19878 | 0.28151 | 0.28148 | 0.28179 | 0.28175 |
| $r^2$ | Mean | 0.04614 | 0.04622 | 0.08123 | 0.08124 | 0.08133 | 0.08132 |
| | SD | 0.07287 | 0.07286 | 0.09796 | 0.09797 | 0.09881 | 0.09882 |
| $D$ | Mean | 0.00154 | 0.00154 | 0.00463 | 0.00463 | 0.00462 | 0.00462 |
| | SD | 0.01197 | 0.01198 | 0.02034 | 0.02035 | 0.02041 | 0.02042 |
| $D'$ | Mean | -0.68313 | -0.68258 | -0.54537 | -0.54544 | -0.54473 | -0.54478 |
| | SD | 0.33979 | 0.33993 | 0.39894 | 0.39878 | 0.39972 | 0.39962 |

| Divergence | Mean | 0.00022 | 0.00070 | 0.00076 | 0.00100 | 0.00075 | 0.00156 |
|---|---|---|---|---|---|---|---|
| | SD | 0.00024 | 0.00822 | 0.00100 | 0.00160 | 0.00099 | 0.00841 |

**Table S4**: Posterior distribution of each parameter inferred by our method. Shown are the posterior probabilities (*prob*) of parameter values presented in bins (*a*, *a*+*bin size*).

| | $f_0$ | $f_1$ | $f_2$ | $f_3$ | $N_{anc}$ | | $N_{cur}$ | | **time** | |
|---|---|---|---|---|---|---|---|---|---|---|
| *a* | *prob* | *prob* | *prob* | *prob* | *a* | *prob* | *a* | *prob* | *a* | *prob* |
| -5 | | | 0.0004 | 0.6829 | 3000 | 0.0003 | 5000 | 0.0000 | 150 | 0.0000 |
| 0 | | 0.0000 | 0.0023 | 0.3094 | 4000 | 0.0022 | 10000 | 0.0000 | 200 | 0.0174 |
| 5 | | 0.0000 | 0.0383 | 0.0068 | 5000 | 0.0331 | 15000 | 0.0001 | 250 | 0.1584 |
| 10 | 0.0000 | 0.0003 | 0.1179 | 0.0009 | 6000 | 0.1699 | 20000 | 0.0007 | 300 | 0.3990 |
| 15 | 0.0000 | 0.0083 | 0.1264 | | 7000 | 0.3718 | 25000 | 0.0061 | 350 | 0.3313 |
| 20 | 0.0007 | 0.0699 | 0.1518 | | 8000 | 0.2895 | 30000 | 0.0558 | 400 | 0.0834 |
| 25 | 0.0006 | 0.1382 | 0.1479 | | 9000 | 0.0979 | 35000 | 0.1597 | 450 | 0.0092 |
| 30 | 0.0043 | 0.1849 | 0.1225 | | 10000 | 0.0258 | 40000 | 0.3501 | 500 | 0.0014 |
| 35 | 0.0391 | 0.1785 | 0.1606 | | 11000 | 0.0061 | 45000 | 0.3347 | 550 | 0.0000 |
| 40 | 0.2341 | 0.1710 | 0.1047 | | 12000 | 0.0023 | 50000 | 0.0854 | 600 | 0.0000 |
| 45 | 0.3636 | 0.1462 | 0.0248 | | 13000 | 0.0006 | 55000 | 0.0074 | | |
| 50 | 0.2229 | 0.0540 | 0.0017 | | 14000 | 0.0005 | 60000 | 0.0000 | | |
| 55 | 0.0918 | 0.0255 | 0.0007 | | 15000 | 0.0002 | 65000 | 0.0000 | | |
| 60 | 0.0335 | 0.0064 | 0.0000 | | 16000 | 0.0000 | | | | |
| 65 | 0.0093 | 0.0123 | 0.0000 | | | | | | | |
| 70 | 0.0001 | 0.0040 | 0.0000 | | | | | | | |
| 75 | | 0.0004 | 0.0000 | | | | | | | |
| 80 | | 0.0000 | | | | | | | | |
| 85 | | | | | | | | | | |
| 90 | | | | | | | | | | |
| 95 | | | | | | | | | | |
| 100 | | | | | | | | | | |

**Table S5**: Distribution of lengths of phastCons elements in the human genome.

| Quantile | 25% | 50% | 75% | 90% | 95% | 97% | 99% |
|---|---|---|---|---|---|---|---|
| Length (bp) | 6 | 10 | 17 | 32 | 51 | 70 | 132 |

**Table S6**: Effects of background selection (shown as $B$ values) at different distances from phastCons elements ($y$) of varying lengths and cross-over rates. The assumed DFE here was one highly skewed towards mildly deleterious mutations such that $f_0 = 0.1;\ f_1 = 0.7;\ f_2 = 0.1;\ f_3 = 0.1$. The mean rate of cross-over is assumed to be $1 \times 10^{-8}$per site per generation.

| | | ***B*** | | |
|---|---|---|---|---|
| **Length (of phastCons element)** | **Recombination rate** | ***y*=1 bp** | ***y*=500 bp** | ***y*=5000 bp** |
| 10 bp | 0.1 × mean | 0.999 | 0.999 | 1.000 |
| | mean | 0.999 | 1.000 | 1.000 |
| | 10 × mean | 0.999 | 1.000 | 1.000 |
| 50 bp | 0.1 × mean | 0.997 | 0.997 | 0.998 |
| | mean | 0.997 | 0.998 | 0.999 |
| | 10 × mean | 0.998 | 0.999 | 1.000 |
| 150 bp | 0.1 × mean | 0.992 | 0.993 | 0.993 |
| | mean | 0.993 | 0.993 | 0.997 |
| | 10 × mean | 0.993 | 0.997 | 1.000 |

**Table S7**: Bias in inference when (1) ancestral alleles are mis-specified in data and (2) mutation rates vary across exons but are assumed to be constant. *To test mis-specified ancestral state*: 1% of all derived singletons were assumed to be falsely polarized and were thus randomly re-assigned to an allele frequency of 99%. *To test both model violations*: Simulated sites were filtered according to the 1000 Genomes Phase 3 data (intergenic sites were filtered to resemble the filtering employed in the 5′ intergenic regions). Inference was performed 50 times and the mean and standard deviation (SD) of those inferred values are presented below.

| | | | $f_0$ | $f_1$ | $f_2$ | $f_3$ | $N_{anc}$ | $N_{cur}$ | time |
|---|---|---|---|---|---|---|---|---|---|
| Equilibrium model | | | | | | | | | |
| True values | | | 0.25 | 0.25 | 0.25 | 0.25 | 20000 | 20000 | 800 |
| Inferred values | Constant mutation rate | Mean | 0.26 | 0.23 | 0.26 | 0.25 | 20035 | 19466 | 619 |
| | | SD | 0.02 | 0.03 | 0.03 | 0.01 | 789 | 937 | 41 |
| | Variable mutation rate | Mean | 0.34 | 0.07 | 0.28 | 0.31 | 21251 | 25836 | 777 |
| | | SD | 0.02 | 0.01 | 0.03 | 0.03 | 1128 | 1669 | 52 |
| | Mis-specified ancestral state | Mean | 0.28 | 0.17 | 0.36 | 0.2 | 20203 | 18891 | 607 |
| | | SD | 0.03 | 0.04 | 0.04 | 0.03 | 850 | 1052 | 42 |
| Growth model | | | | | | | | | |
| True values | | | 0.25 | 0.25 | 0.25 | 0.25 | 20000 | 40000 | 800 |
| Inferred values | Constant mutation rate | Mean | 0.21 | 0.23 | 0.26 | 0.30 | 20127 | 30911 | 752 |
| | | SD | 0.02 | 0.02 | 0.03 | 0.02 | 427 | 2282 | 47 |
| | Variable mutation rate | Mean | 0.35 | 0.18 | 0.31 | 0.17 | 18409 | 47759 | 809 |
| | | SD | 0.01 | 0.02 | 0.01 | 0.01 | 726 | 1087 | 15 |
| | Mis-specified ancestral state | Mean | 0.25 | 0.16 | 0.35 | 0.23 | 20200 | 29655 | 733 |
| | | SD | 0.03 | 0.03 | 0.04 | 0.02 | 754 | 2302 | 72 |
| Decline model | | | | | | | | | |
| True values | | | 0.25 | 0.25 | 0.25 | 0.25 | 20000 | 10000 | 800 |
| Inferred values | Constant mutation rate | Mean | 0.23 | 0.29 | 0.21 | 0.27 | 19694 | 9645 | 416 |
| | | SD | 0.00 | 0.01 | 0.02 | 0.02 | 195 | 571 | 8 |

| | | | | | | | | | |
|---|---|---|---|---|---|---|---|---|---|
| | Variable mutation rate | Mean | 0.30 | 0.10 | 0.43 | 0.17 | 20414 | 6904 | 297 |
| | | SD | 0.01 | 0.01 | 0.01 | 0.01 | 348 | 620 | 20 |
| | Mis-specified ancestral state | Mean | 0.30 | 0.14 | 0.32 | 0.24 | 20462 | 10957 | 424 |
| | | SD | 0.01 | 0.01 | 0.01 | 0.02 | 236 | 825 | 8 |

**Table S8**: URLs of downloaded data.

| Data | Link |
|---|---|
| Gene annotations (GRCh37) | https://www.ncbi.nlm.nih.gov/projects/genome/guide/human/ |
| phastConsElements100way | https://genome.ucsc.edu/cgi-bin/hgTables?hgsid=1503445671_LLYCngZX0ohcfgUJahGGhVofwA2j&clade=mammal&org=Human&db=hg19&hgta_group=compGeno&hgta_track=cons100way&hgta_table=phastConsElements100way&hgta_regionType=genome&position=chrX%3A15%2C578%2C261-15%2C621%2C068&hgta_outputType=wigData&hgta_outFileName=strictAccs_1000G_all |
| sno/miRNAs | https://genome.ucsc.edu/cgi-bin/hgTables?hgsid=1503445671_LLYCngZX0ohcfgUJahGGhVofwA2j&clade=mammal&org=Human&db=hg19&hgta_group=genes&hgta_track=wgRna&hgta_table=0&hgta_regionType=genome&position=chrX%3A15%2C578%2C261-15%2C621%2C068&hgta_outputType=primaryTable&hgta_outFileName=strictAccs_1000G_all (last updated 2010-09-20) |
| Recombination rates | https://genome.ucsc.edu/cgi-bin/hgTables?hgsid=1267722847_s57UAaaSF8GUOKQv04APCh6JAZY5&clade=mammal&org=Human&db=hg19&hgta_group=map&hgta_track=decodeRmap&hgta_table=hapMapRelease24YRIRecombMap&hgta_regionType=genome&position=chr21%3A33%2C031%2C597-33%2C041%2C570&hgta_outputType=wigData&hgta_outFileName= |
| Strict accessibility files | https://genome.ucsc.edu/cgi-bin/hgTables?hgsid=1444498527_jTTaja4hABz4K5EVHVnhMN9iK4TR&clade=mammal&org=Human&db=hg19&hgta_group=varRep&hgta_track=tgpPhase3Accessibility&hgta_table=tgpPhase3AccessibilityStrictCriteria&hgta_regionType=genome&position=chrX%3A15%2C578%2C261-15%2C621%2C068&hgta_outputType=primaryTable&hgta_outFileName=strictAccs_1000G_all |
| Variants from 1000 Genomes Phase 3 | http://hgdownload.soe.ucsc.edu/gbdb/hg19/1000Genomes/phase3/ |
| Hg19 reference genome | http://hgdownload.soe.ucsc.edu/goldenPath/hg19/chromosomes/ |
| Ancestral alleles | ftp://ftp.ensembl.org/pub/release-74/fasta/ancestral_alleles/homo_sapiens_ancestor_GRCh37_e71.tar.bz2 |
| Mutation rate maps | https://www.nlgenome.nl/menu/main/app-download |

**Table S9**: Individuals from the 1000 Genomes Phase 3 YRI individuals used for inference.

| **number** | **sample** | **pop** | **sex** |
|---|---|---|---|
| 1 | NA18488 | YRI | female |
| 2 | NA18489 | YRI | female |
| 3 | NA18499 | YRI | female |
| 4 | NA18502 | YRI | female |
| 5 | NA18505 | YRI | female |
| 6 | NA18508 | YRI | female |
| 7 | NA18511 | YRI | female |
| 8 | NA18517 | YRI | female |
| 9 | NA18520 | YRI | female |
| 10 | NA18523 | YRI | female |
| 11 | NA18858 | YRI | female |
| 12 | NA18861 | YRI | female |
| 13 | NA18864 | YRI | female |
| 14 | NA18867 | YRI | female |
| 15 | NA18870 | YRI | female |
| 16 | NA18873 | YRI | female |
| 17 | NA18876 | YRI | female |
| 18 | NA18878 | YRI | female |
| 19 | NA18881 | YRI | female |
| 20 | NA18907 | YRI | female |
| 21 | NA18909 | YRI | female |
| 22 | NA18912 | YRI | female |
| 23 | NA18916 | YRI | female |
| 24 | NA18924 | YRI | female |
| 25 | NA18933 | YRI | female |
| 1 | NA18486 | YRI | male |
| 2 | NA18498 | YRI | male |
| 3 | NA18501 | YRI | male |
| 4 | NA18504 | YRI | male |
| 5 | NA18507 | YRI | male |
| 6 | NA18510 | YRI | male |
| 7 | NA18516 | YRI | male |
| 8 | NA18519 | YRI | male |
| 9 | NA18522 | YRI | male |
| 10 | NA18853 | YRI | male |
| 11 | NA18856 | YRI | male |
| 12 | NA18865 | YRI | male |
| 13 | NA18868 | YRI | male |
| 14 | NA18871 | YRI | male |
| 15 | NA18874 | YRI | male |
| 16 | NA18877 | YRI | male |
| 17 | NA18879 | YRI | male |
| 18 | NA18908 | YRI | male |

| | | | |
|---|---|---|---|
| 19 | NA18910 | YRI | male |
| 20 | NA18915 | YRI | male |
| 21 | NA18917 | YRI | male |
| 22 | NA18923 | YRI | male |
| 23 | NA18934 | YRI | male |
| 24 | NA19092 | YRI | male |
| 25 | NA19096 | YRI | male |

**Table S10**: Statistics obtained using all individuals from the YRI population (post filtering).

| | | 5′ less linked | 5′ linked | exonic | 3′ linked | 3′ less linked |
|---|---|---|---|---|---|---|
| $\pi$ | Mean | 0.00091 | 0.00087 | 0.00073 | 0.00094 | 0.00098 |
| | SD | 0.00106 | 0.00078 | 0.00053 | 0.00101 | 0.00107 |
| $\theta_W$ | Mean | 0.00114 | 0.00113 | 0.00103 | 0.00113 | 0.00120 |
| | SD | 0.00083 | 0.00072 | 0.00054 | 0.00076 | 0.00104 |
| $\theta_H$ | Mean | 0.00086 | 0.00081 | 0.00068 | 0.00095 | 0.00099 |
| | SD | 0.00134 | 0.00116 | 0.00073 | 0.00153 | 0.00170 |
| $H'$ | Mean | 0.01196 | 0.02367 | 0.05312 | -0.03075 | -0.01562 |
| | SD | 0.83960 | 0.79121 | 0.68490 | 0.99036 | 0.92945 |
| Tajima's $D$ | Mean | -0.46977 | -0.49783 | -0.79460 | -0.44045 | -0.42656 |
| | SD | 0.85756 | 0.80704 | 0.71098 | 0.81487 | 0.83339 |
| Singleton density | Mean | 0.00168 | 0.00159 | 0.00162 | 0.00147 | 0.00146 |
| | SD | 0.00209 | 0.00169 | 0.00098 | 0.00166 | 0.00170 |
| Haplotype diversity | Mean | 0.49875 | 0.51198 | 0.68764 | 0.51457 | 0.51045 |
| | SD | 0.30316 | 0.29983 | 0.21594 | 0.30976 | 0.29634 |
| $r^2$ | Mean | 0.06821 | 0.06477 | 0.05194 | 0.06475 | 0.06875 |
| | SD | 0.10622 | 0.07743 | 0.04243 | 0.08376 | 0.09427 |
| $D$ | Mean | 0.00462 | 0.00327 | 0.00229 | 0.00385 | 0.00501 |
| | SD | 0.01572 | 0.01134 | 0.00576 | 0.01545 | 0.01966 |
| $D'$ | Mean | -0.65663 | -0.65836 | -0.69276 | -0.62330 | -0.62955 |
| | SD | 0.34114 | 0.32467 | 0.21832 | 0.39850 | 0.36866 |
| Divergence | Mean | 0.00563 | 0.00540 | 0.00449 | 0.00545 | 0.00580 |
| | SD | 0.00616 | 0.00488 | 0.00616 | 0.00437 | 0.00585 |

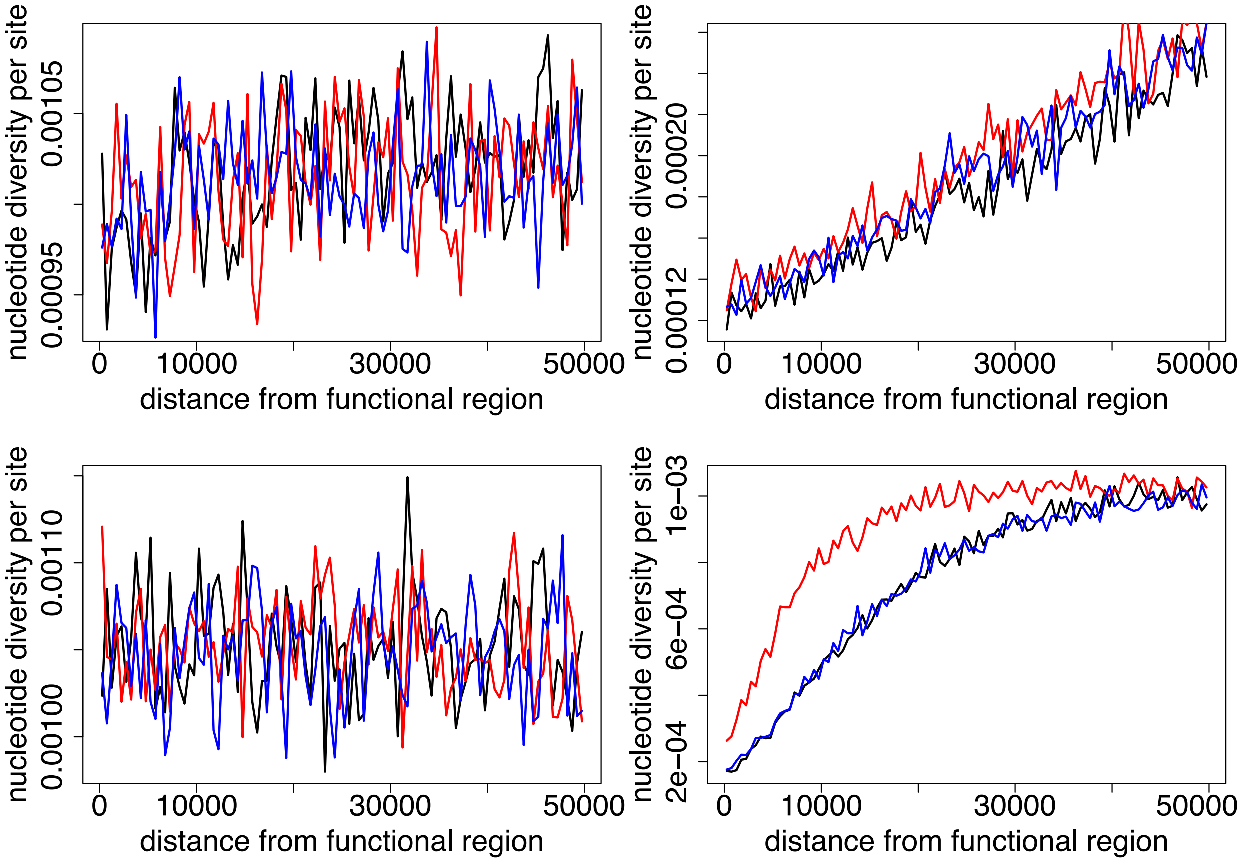


**Figure S1**: Effect of gene conversion on the recovery of nucleotide diversity around functional elements (of 4 kb in length) due to hitchhiking effects. The left panel displays the recovery due to only background selection and right panel shows the recovery in the presence of both background selection and selective sweeps. The cross-over rate is the mean rate (of $10^{-8}$ per site/gen) in the top panel and is 10-fold the mean rate in the bottom panel. Shown are cases for no gene conversion (black lines), a fixed rate of gene conversion of $4.72 \times 10^{-8}$ per site/gen (blue lines) and the case where rates of gene conversion are 5-fold that of the cross-over rate (red lines). Background selection effects were generated by the DFE ($f_0 = 0.22, f_1 = 0.27, f_2 = 0.13, f_3 = 0.38$), while sweep effects were generated by recurrent positive selection such that 1% of all new mutations were beneficial with exponentially distributed selective effects of mean $2N_es$=500. Simulations were performed assuming a Wright-Fisher population under equilibrium with $N_e = 23{,}222$ and mutation rate = $1.25 \times 10^{-8}$ per site/gen. Simulations were scaled down by a factor of 10, and 1000 replicates were simulated with the above plots displaying the mean values of statistics across the replicates.

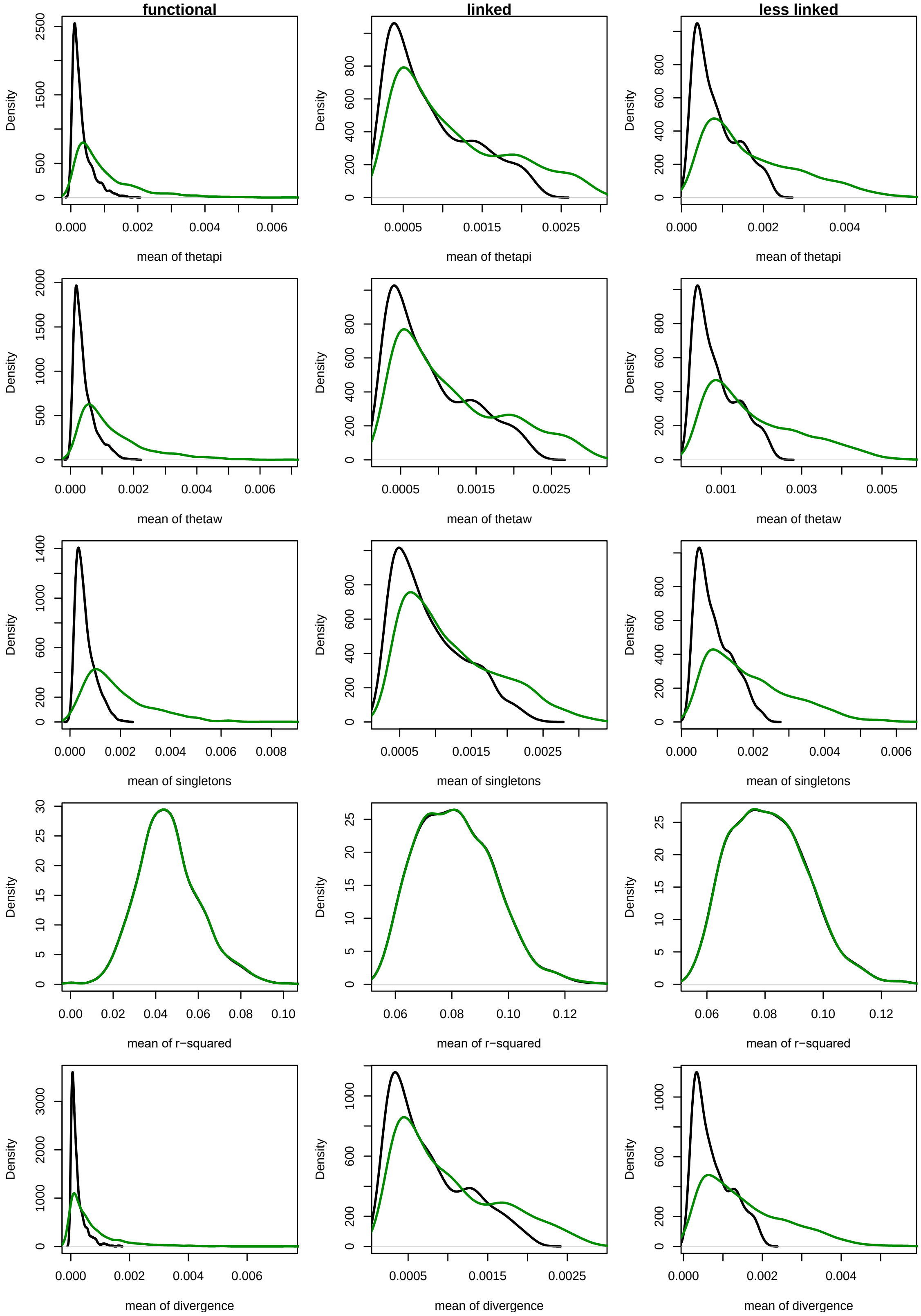


**Figure S2**: Distribution of mean of statistics (across the 465 exons) shown for the first 1000 simulated parameter combinations when no filtering was employed (in black) vs. when sites were filtered as in the empirical data (in green).

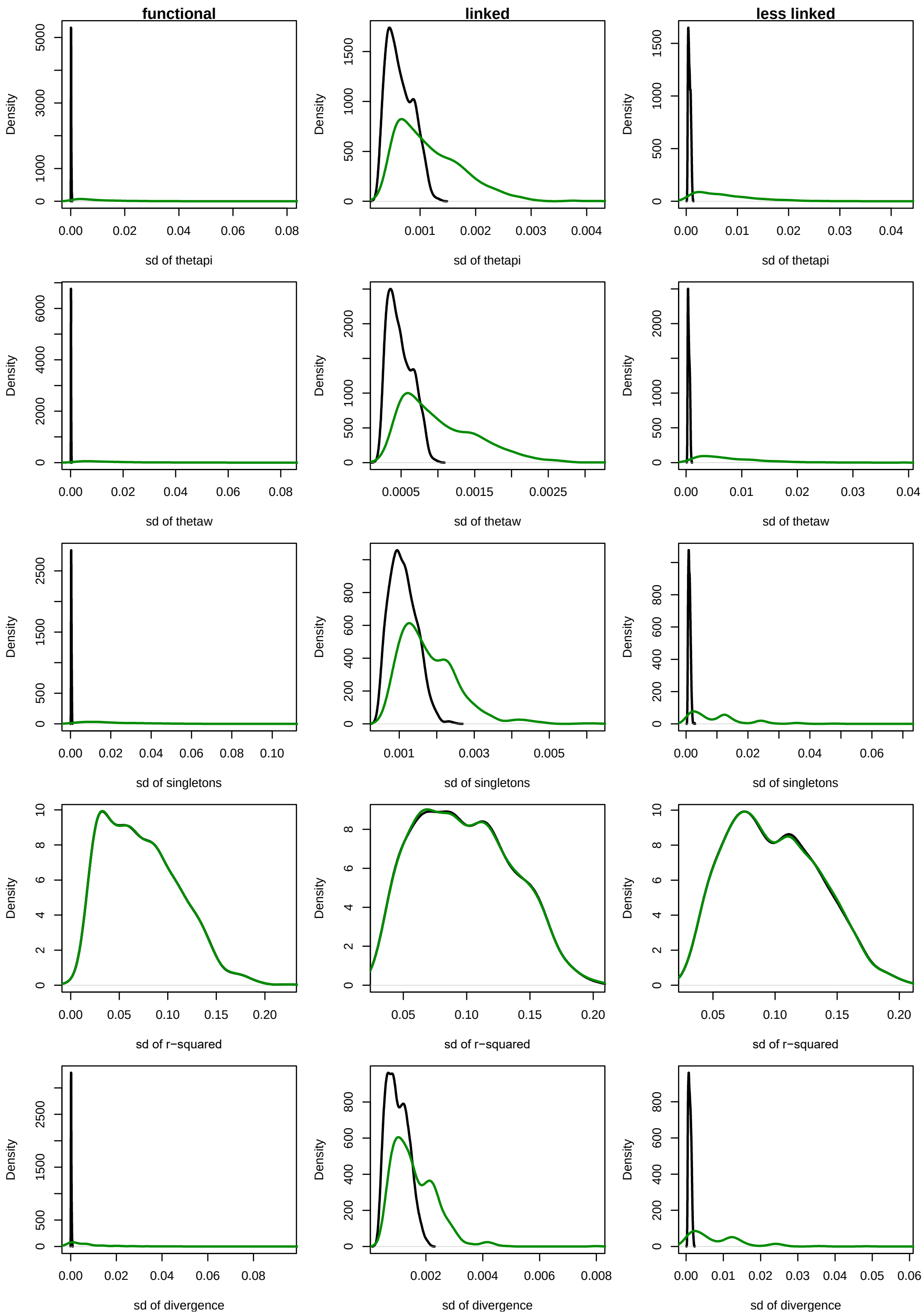


**Figure S3**: Distribution of standard deviation (SD) of statistics (across the 465 exons) shown for the first 1000 simulated parameter combinations when no filtering was employed (in black) vs. when sites were filtered as in the empirical data (in green).

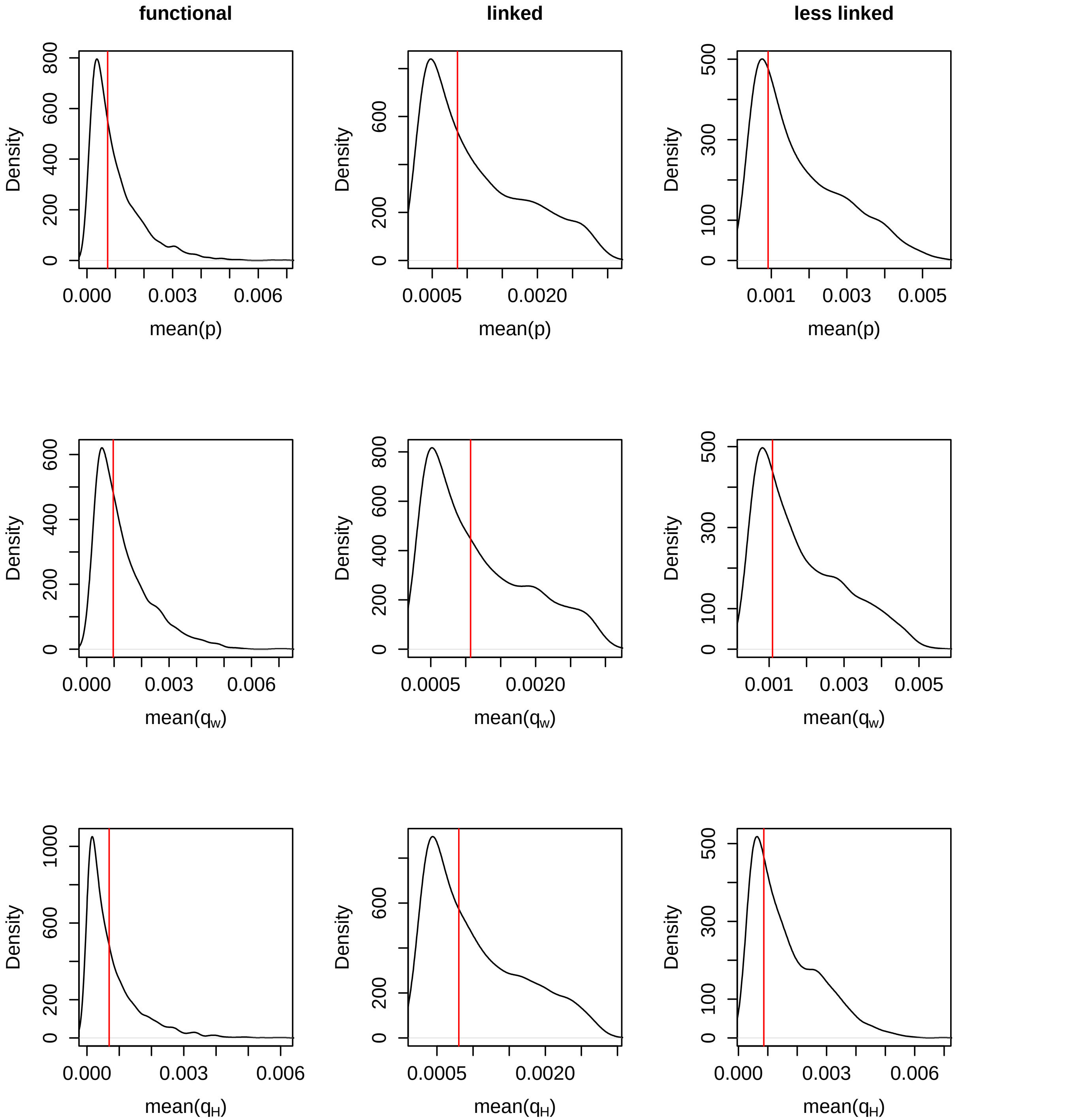


**Figure S4**: Distribution of mean of $\pi$, $\theta_W$, $\theta_H$ across the 465 exons in each of the three windows: “functional”, “linked”, and “less linked” - shown by the black line for all parameter combinations simulated for the purpose of employing ABC in the current study. The red line represents the mean of the same statistics across the 465 exons (and the 5′ intergenic region) observed empirically in the 25 individuals of the YRI population.

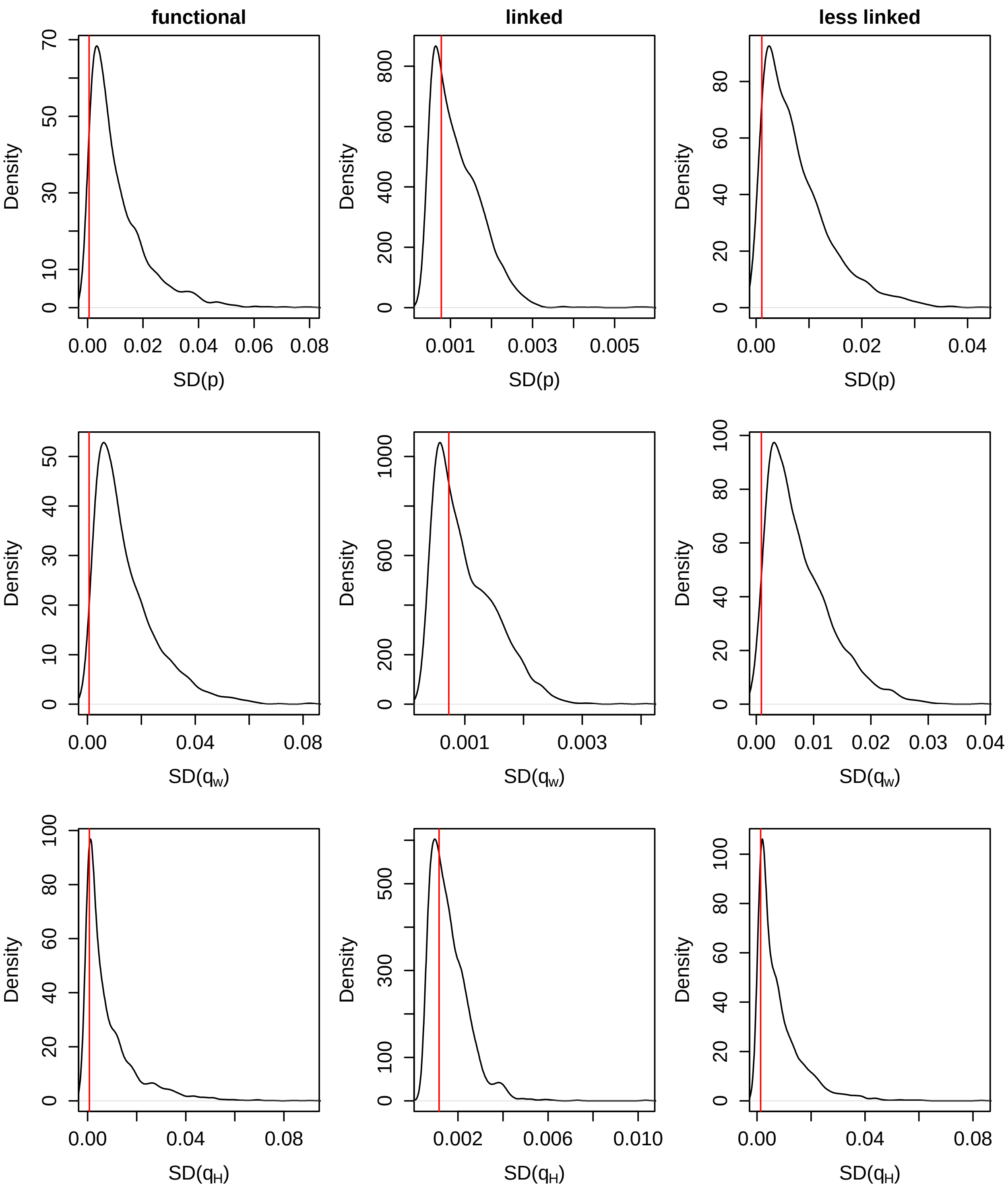


**Figure S5**: Distribution of the variance of $\pi$, $\theta_W$, $\theta_H$ across the 465 exons in each of the three windows: “functional”, “linked”, and “less linked” - shown by the black line for all parameter combinations simulated for the purpose of employing ABC in the current study. The red line represents the mean of the same statistics across the 465 exons (and the 5′ intergenic region) observed empirically in the 50 individuals of the YRI population.

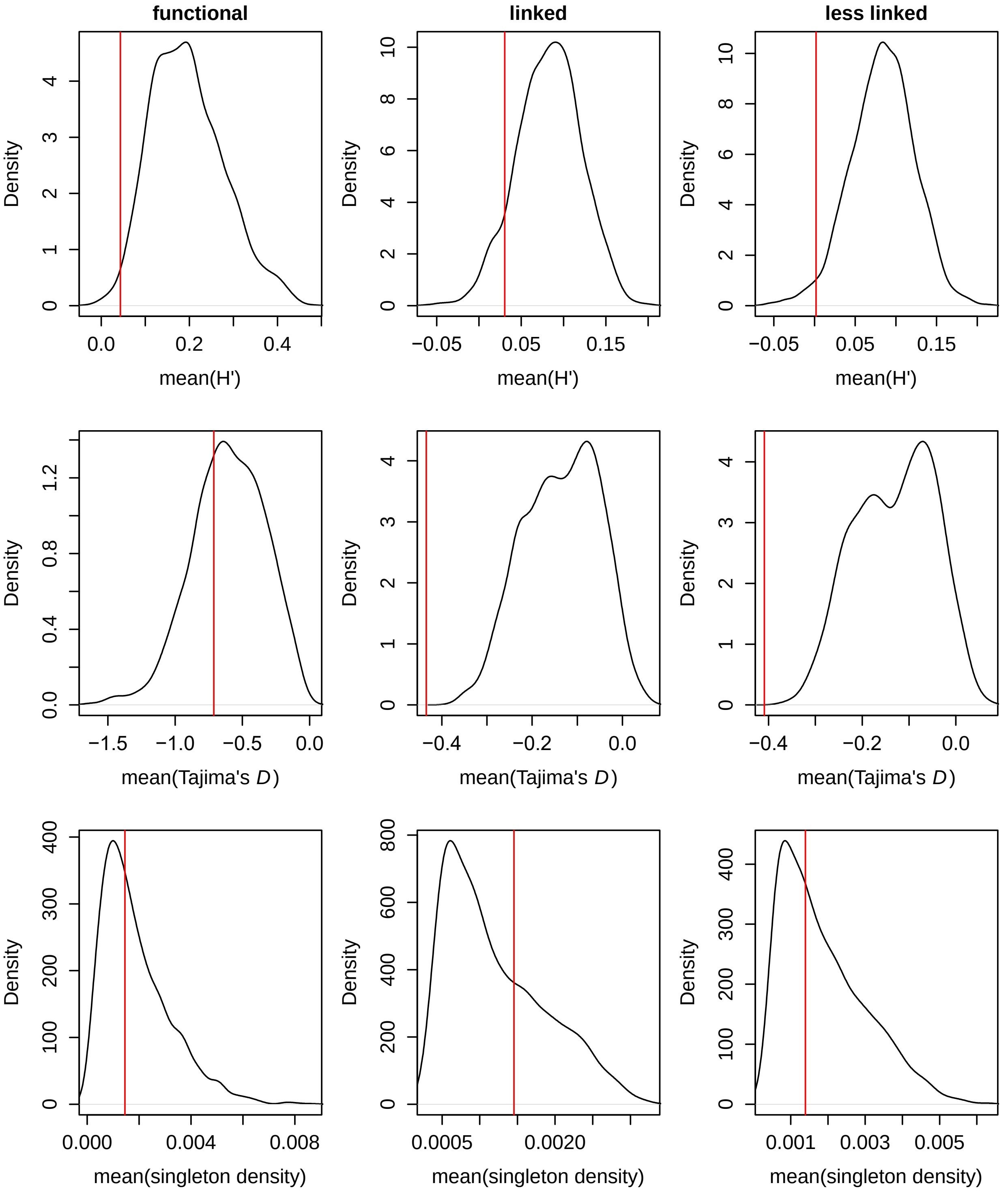


**Figure S6**: Distribution of mean of $H'$, Tajima's $D$, and singleton density across the 465 exons in each of the three windows: "functional", "linked", and "less linked" - shown by the black line for all parameter combinations simulated for the purpose of employing ABC in the current study. The red line represents the mean of the same statistics across the 465 exons (and the 5′ intergenic region) observed empirically in the 50 individuals of the YRI population.

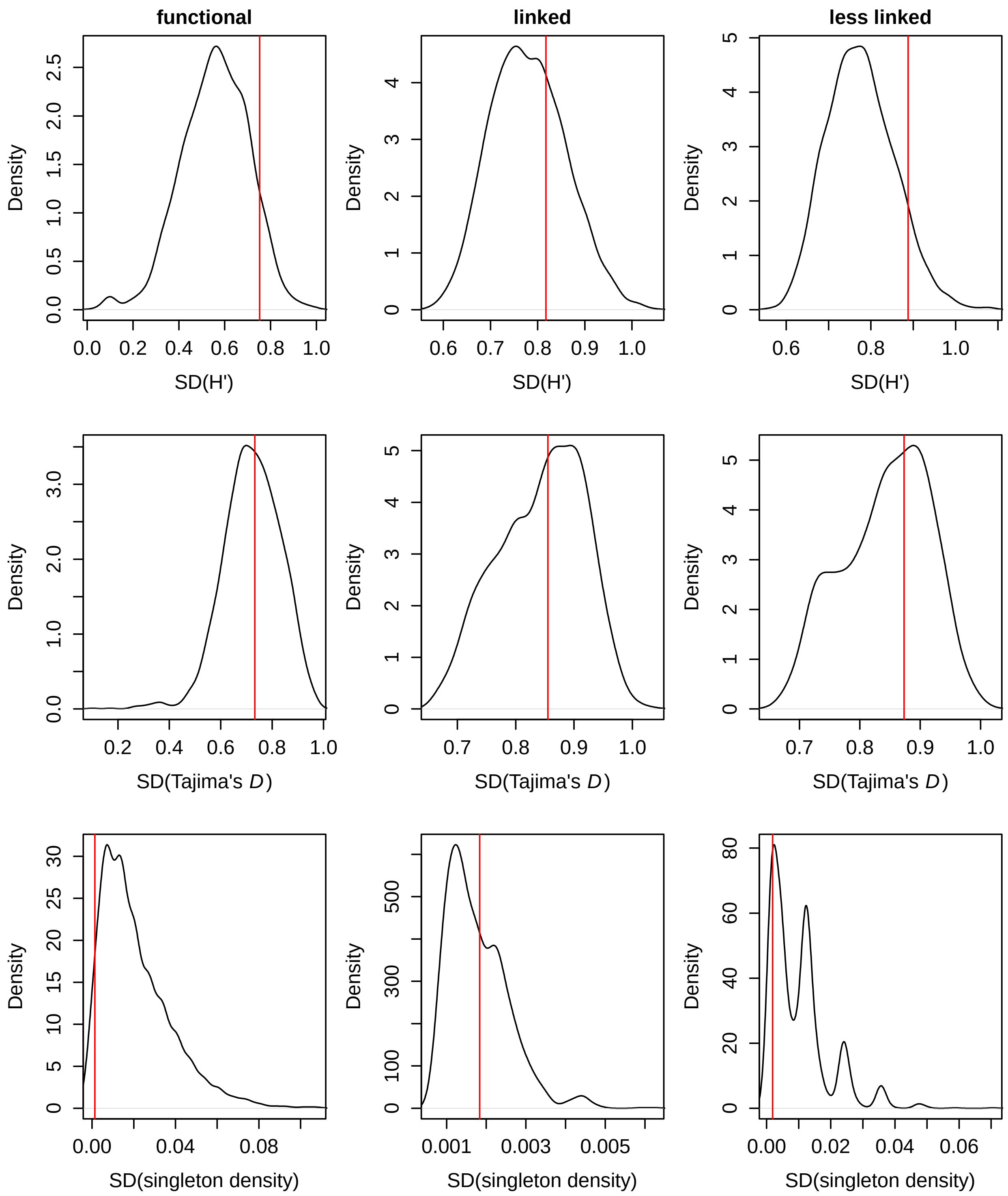


**Figure S7**: Distribution of variance of $H'$, Tajima's $D$, and singleton density across the 465 exons in each of the three windows: "functional", "linked", and "less linked" - shown by the black line for all parameter combinations simulated for the purpose of employing ABC in the current study. The red line represents the mean of the same statistics across the 465 exons (and the 5′ intergenic region) observed empirically in the 50 individuals of the YRI population.

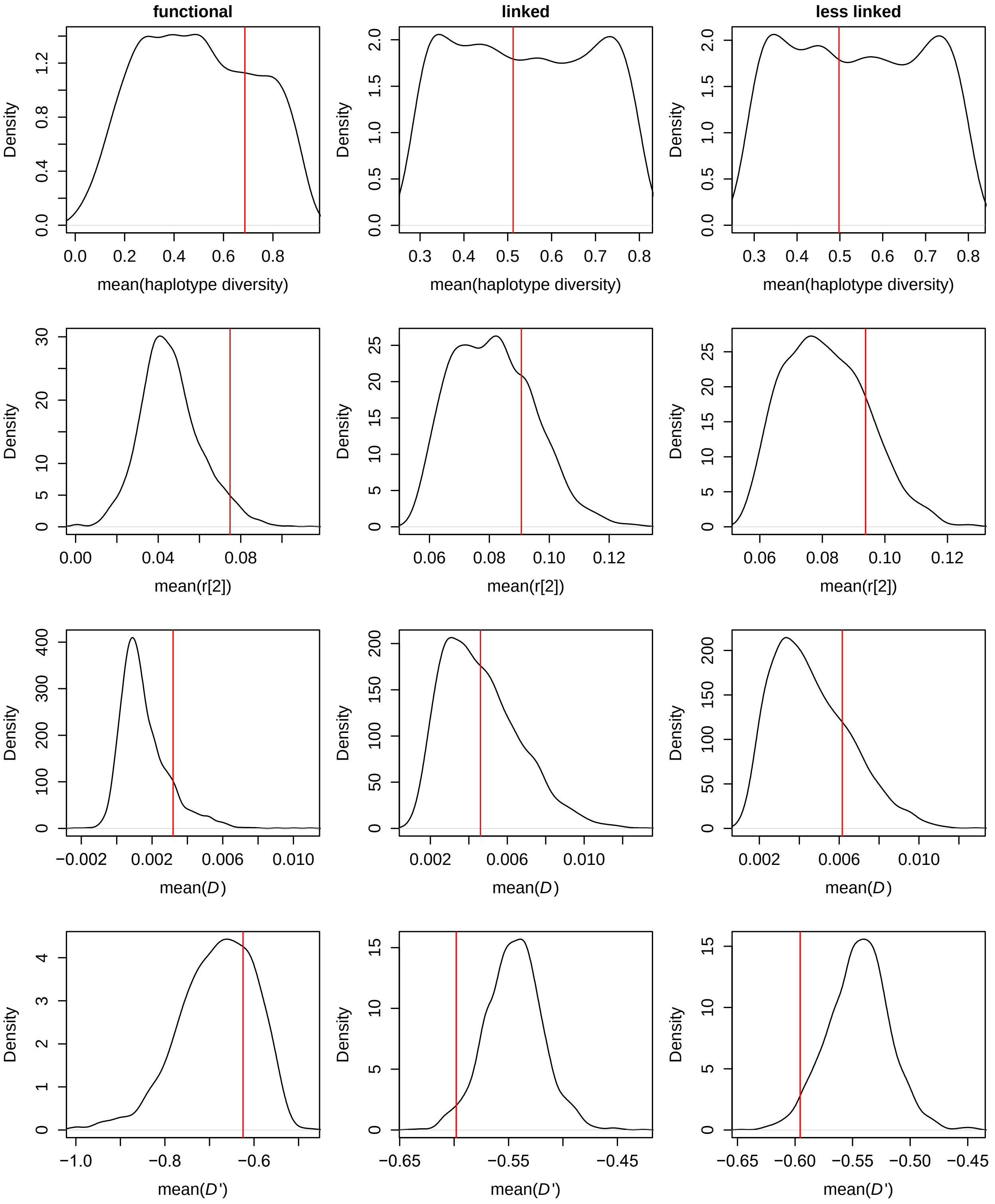


**Figure S8**: Distribution of mean of statistics summarizing LD patterns across the 465 exons in each of the three windows: “functional”, “linked”, and “less linked” - shown by the black line for all parameter combinations simulated for the purpose of employing ABC in the current study. The red line represents the mean of the same statistics across the 465 exons (and the 5′ intergenic region) observed empirically in the 50 individuals of the YRI population.

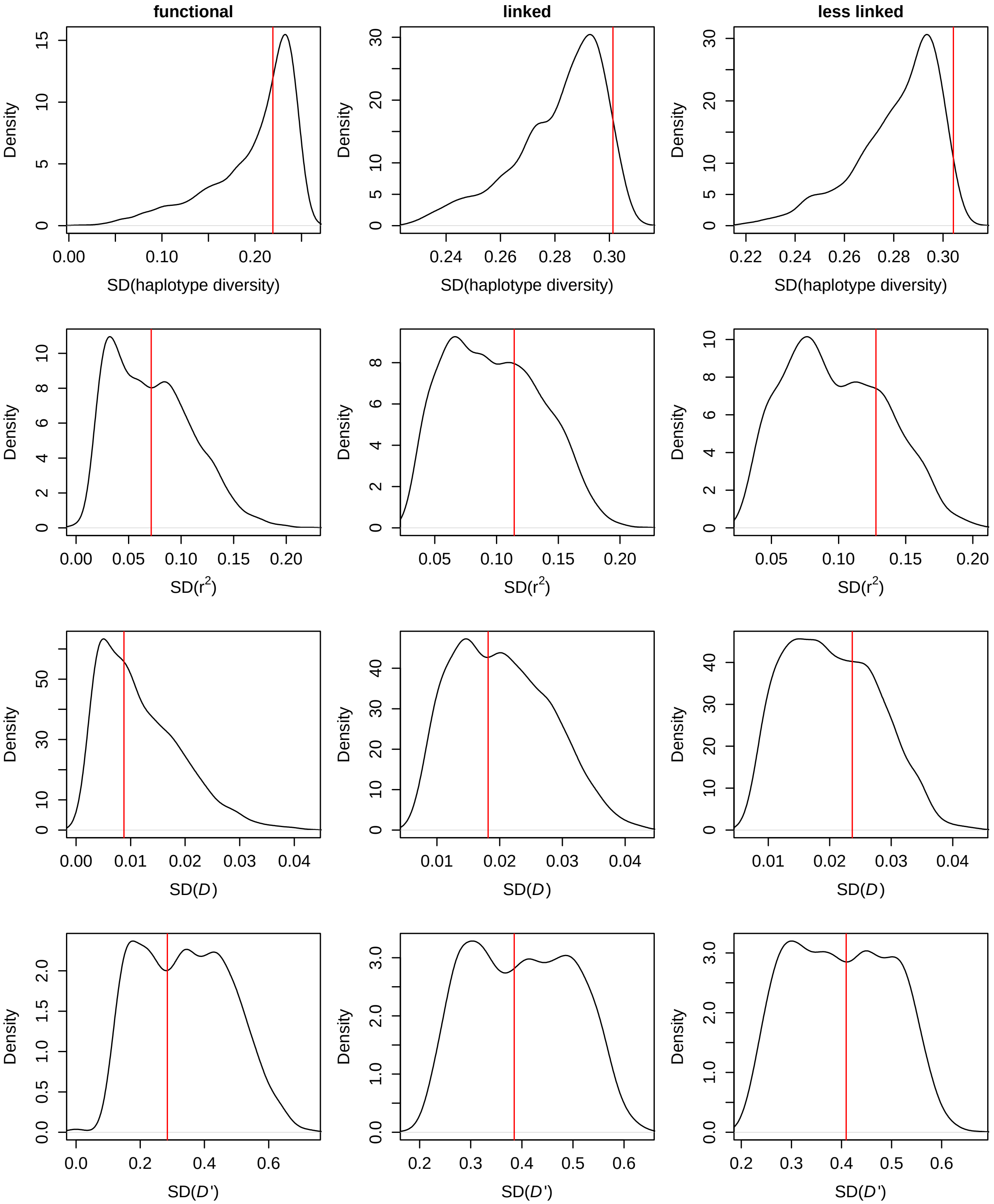


**Figure S9**: Distribution of variance of statistics summarizing LD patterns across the 465 exons in each of the three windows: “functional”, “linked”, and “less linked” - shown by the black line for all parameter combinations simulated for the purpose of employing ABC in the current study. The red line represents the mean of the same statistics across the 465 exons (and the 5′ intergenic region) observed empirically in the 50 individuals of the YRI population.

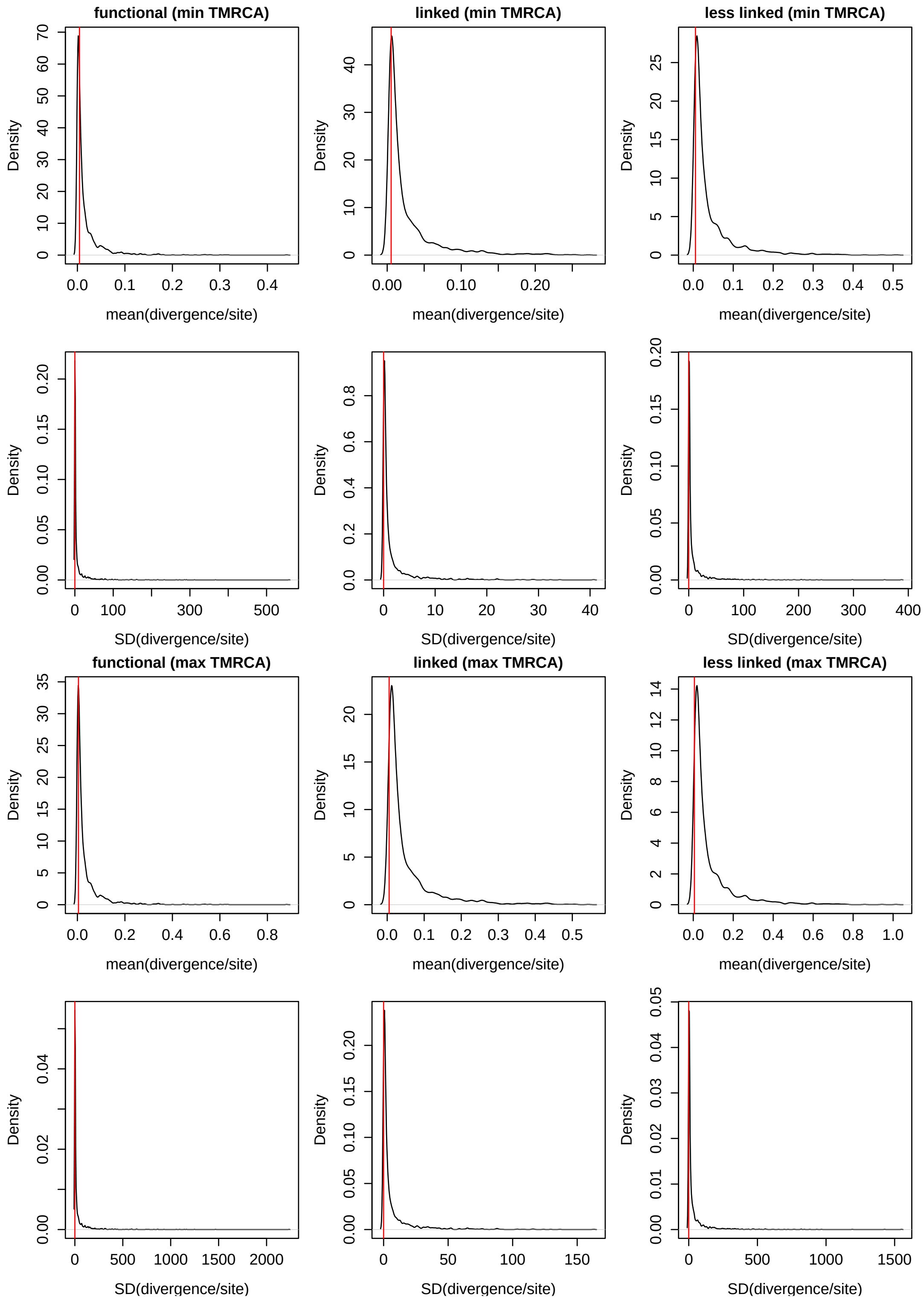


**Figure S10**: Distribution of mean and variance of divergence per site across the 465 exons in each of the three windows: “functional”, “linked”, and “less linked” - shown by the black line for all parameter combinations simulated for the purpose of employing ABC in the current study. The red line represents the mean of the same statistics across the 465 exons (and the 5′ intergenic region) observed empirically in the 50 individuals of the YRI population.

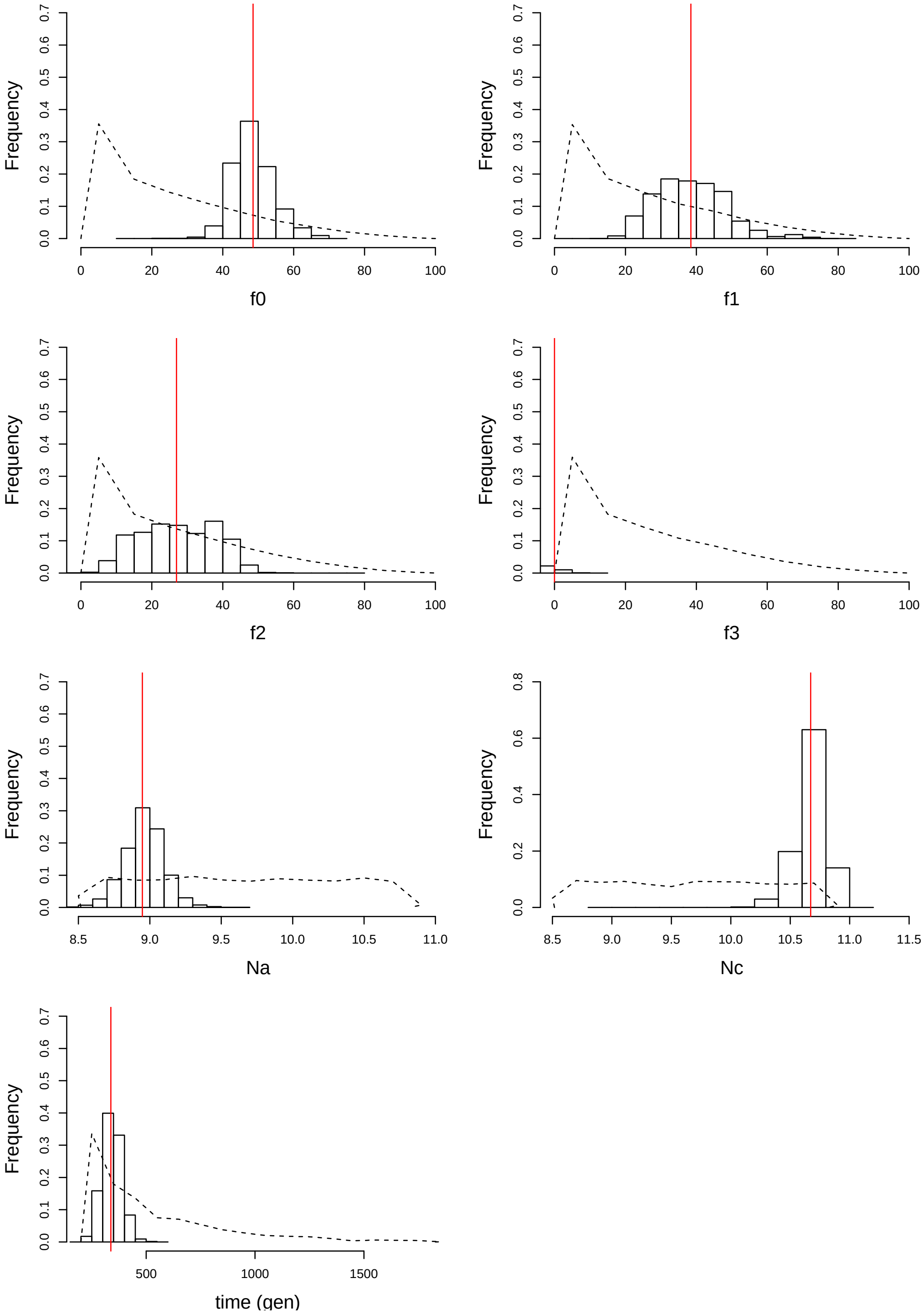


**Figure S11**: Posterior distributions of parameters obtained when performing joint inference of demography and selection. The black dashed line shows the prior distribution, the black solid lines represent the posterior distributions, and the red line shows the final point estimate. Inferences were performed 50 times and the posterior distributions show the distribution of all inferences and the point estimate is calculated as the mean of weighted medians of posterior estimates. The ancestral and current sizes are shown as $\ln{(N_a)}$ and $\ln{(N_c)}$ as their priors were sampled from a log uniform distribution.

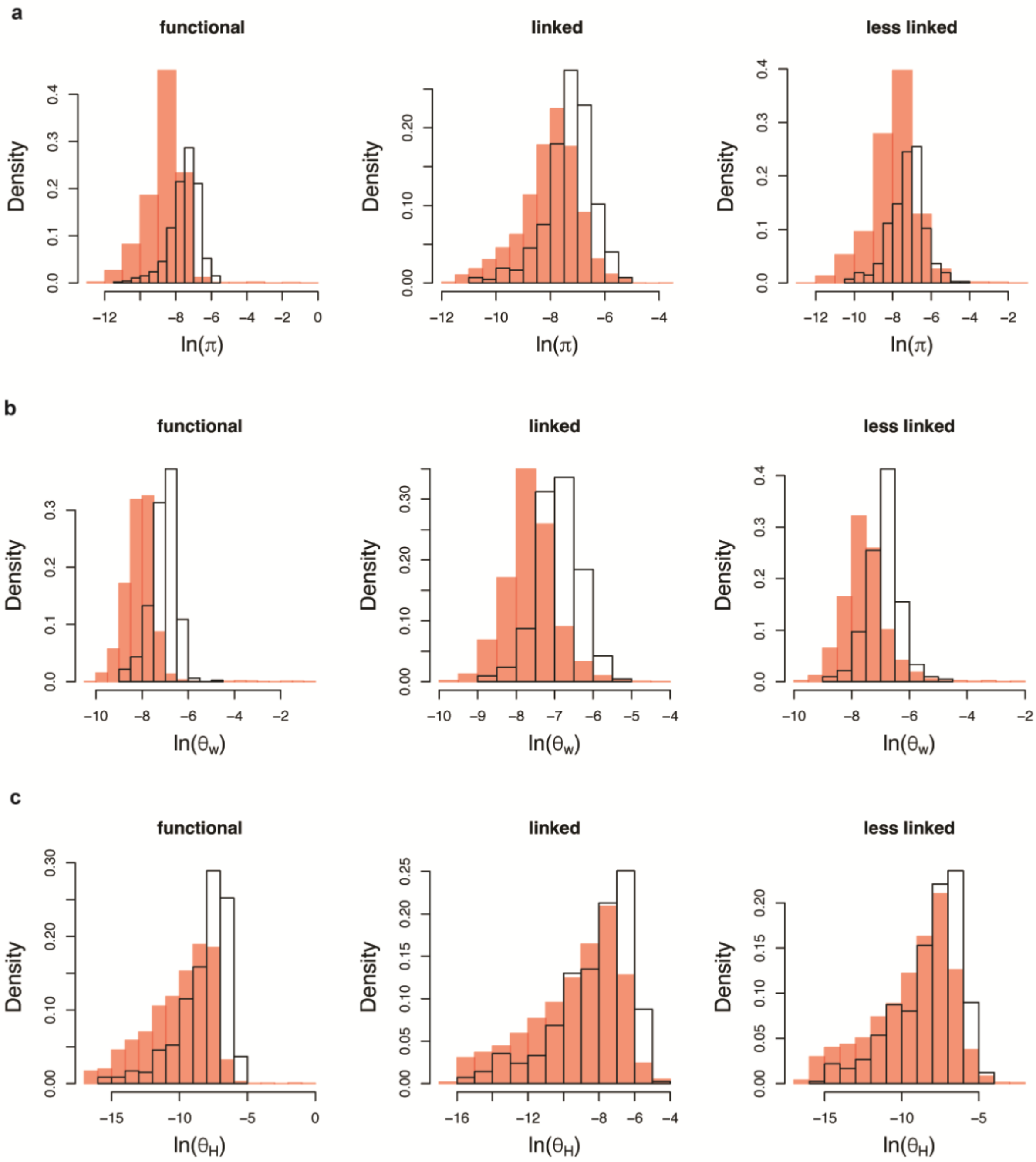


**Figure S12**: Fit of the best model inferred by our method to the empirical data, as shown by the distribution of (a) nucleotide diversity, (b) Watterson's $\theta$, and (c) $\theta_H$, across the 465 exons, for each of the three windows separately – functional, linked, and less linked intergenic regions. The simulated best model (with 10 replicates) is shown in red, while the observed empirical distributions of the same statistics in the YRI population are shown in the white distributions.

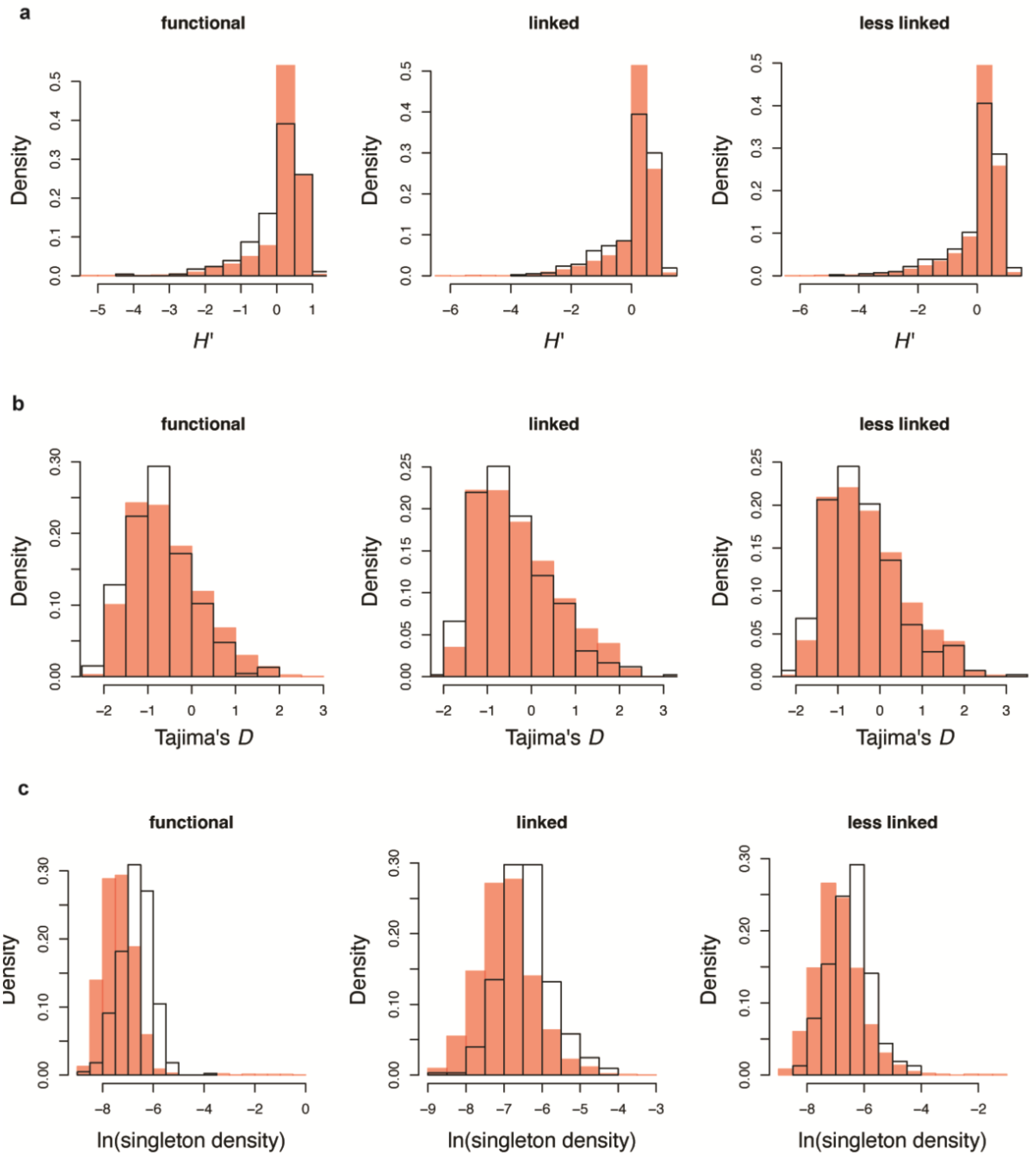


**Figure S13**: Fit of the best model inferred by our method to the empirical data, as shown by the distribution of (a) *H′*, (b) Tajima's *D*, and (c) singleton density, across the 465 exons, for each of the three windows separately: functional, linked, and less linked intergenic regions. The simulated best model (with 10 replicates) is shown in red, while the observed empirical distributions of the same statistics in the YRI population are shown in the white distributions.

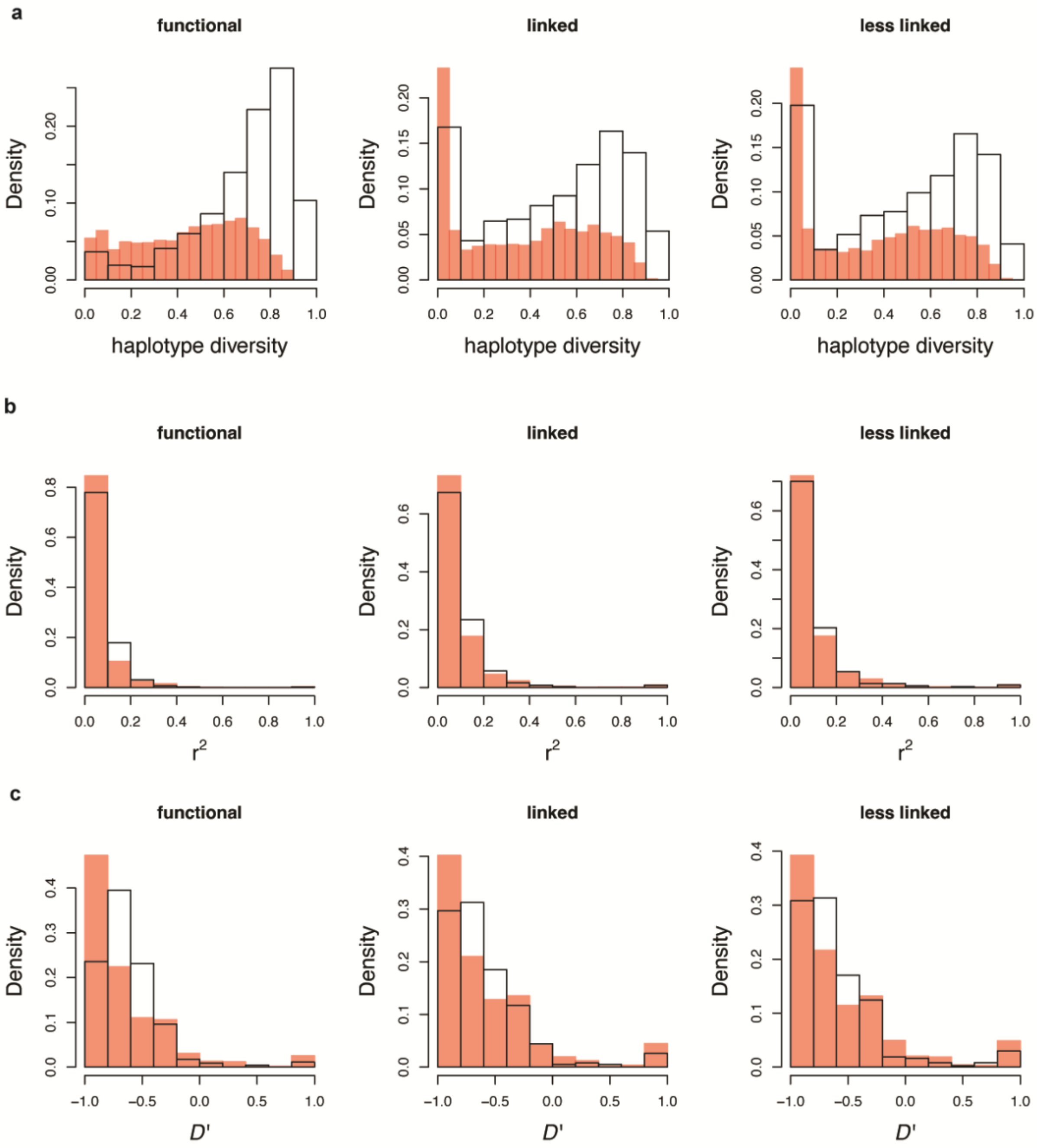


**Figure S14**: Fit of the best model inferred by our method to the empirical data, as shown by the distribution of (a) haplotype diversity, (b) $r^2$, and (c) $D'$, across the 465 exons, for each of the three windows separately: functional, linked, and less linked intergenic regions. The simulated best model (with 10 replicates) is shown in red, while the observed empirical distributions of the same statistics in the YRI population are shown in the white distributions.

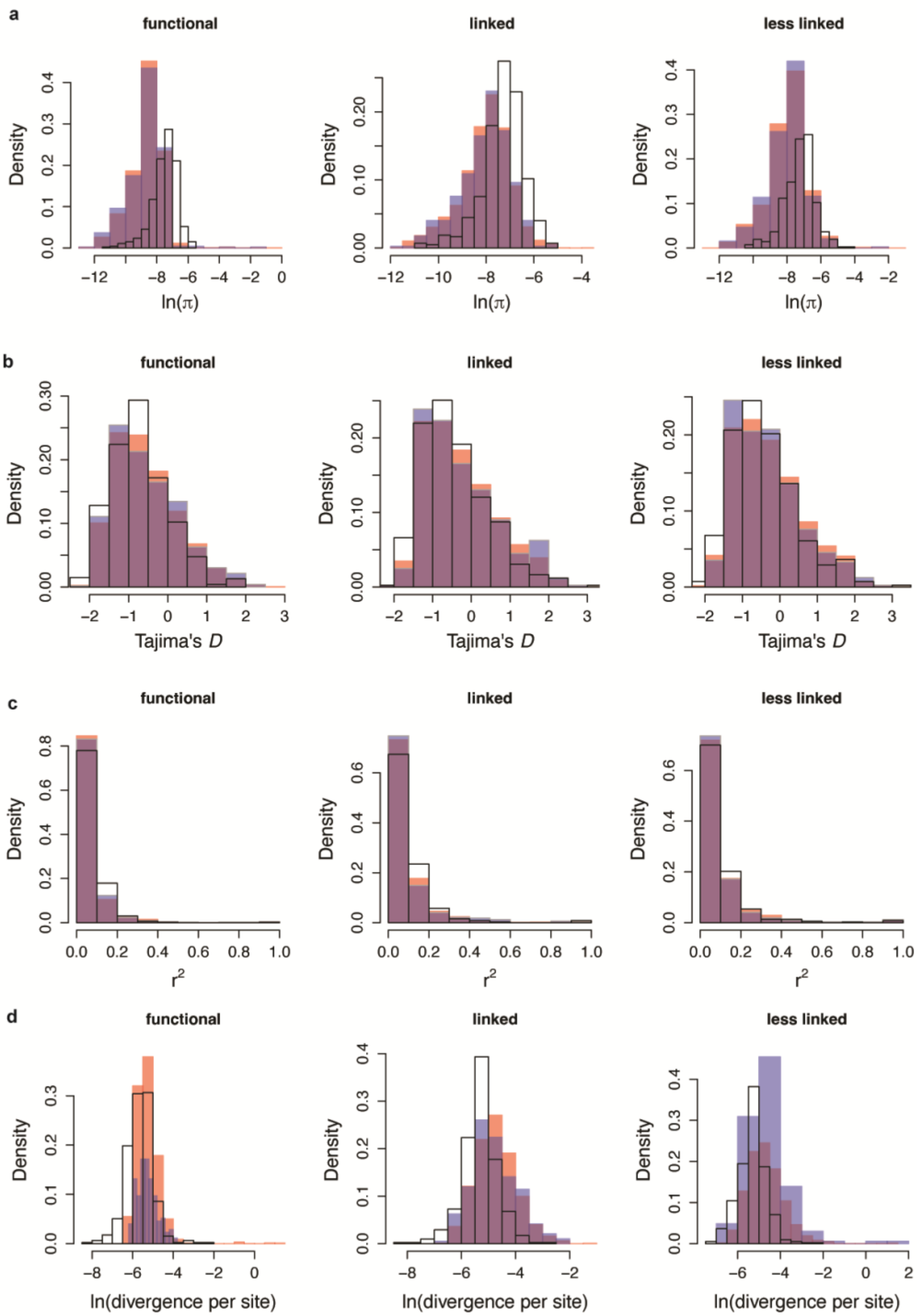


**Figure S15**: Fit of the estimated best model to the empirical data in the presence of weak ($E[2N_e s_b] = 10$) and infrequent ($f_{pos} = 0.1\%$) positive selection. Distribution of (a) nucleotide diversity, (b) Tajima's $D$, (c) $r^2$, and (d) divergence per site across the 465 exons, for each of the three windows separately: functional, linked, and less linked intergenic regions. The best model is depicted in red, the best model with positive selection in blue, and their overlap in purple. The distribution of the empirical data is shown in the white distributions.

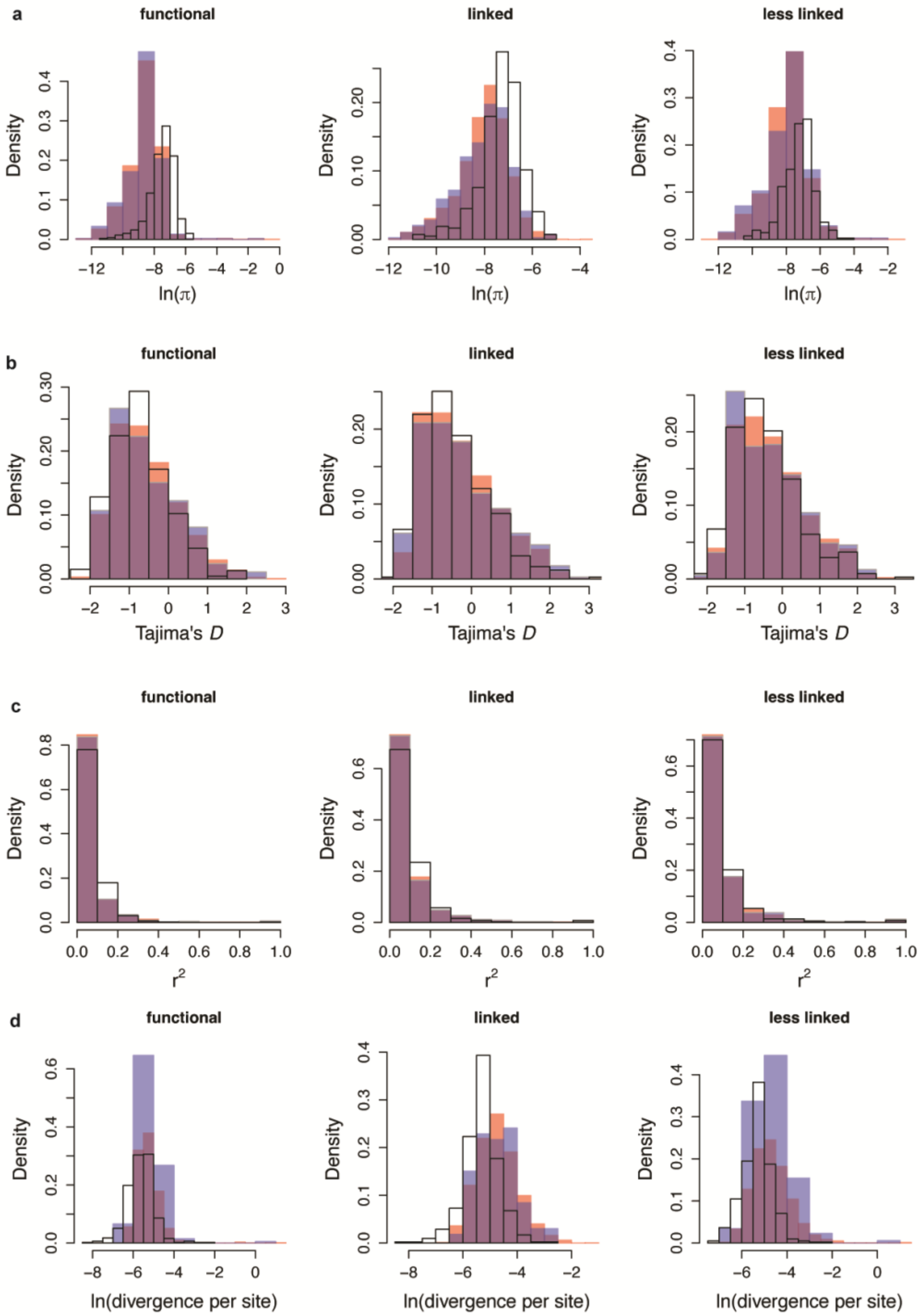


**Figure S16**: Fit of the estimated best model to the empirical data in the presence of moderately strong ($E[2N_e s_b] = 100$) and infrequent ($f_{pos} = 0.1\%$) positive selection. Distribution of (a) nucleotide diversity, (b) Tajima's $D$, (c) $r^2$, and (d) divergence per site across the 465 exons, for each of the three windows separately: functional, linked, and less linked intergenic regions. The best model is depicted in red, the best model with positive selection in blue, and their overlap in purple. The distribution of the empirical data is shown in the white distributions.

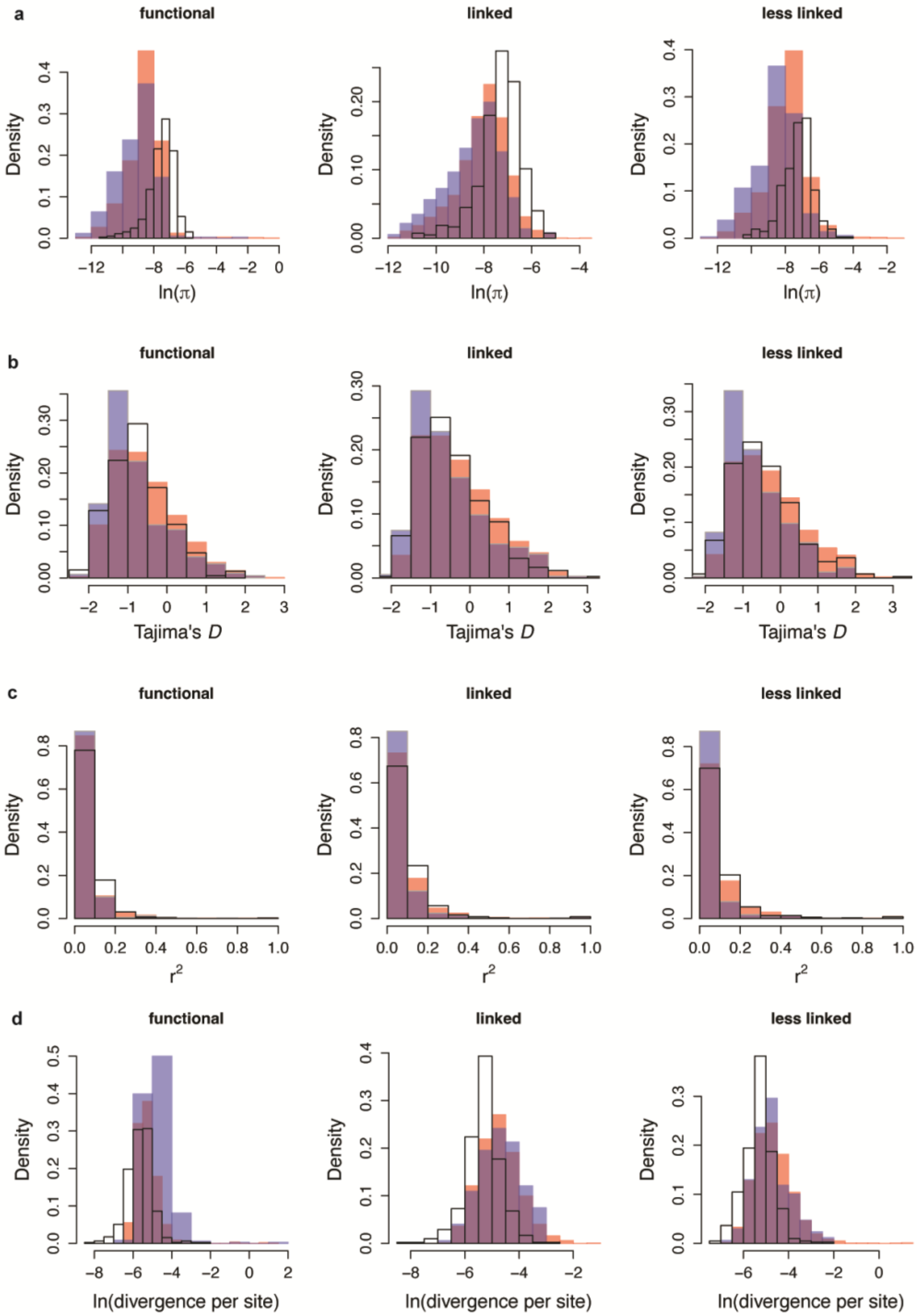


**Figure S17**: Fit of the estimated best model to the empirical data in the presence of strong ($E[2N_e s_b] = 1000$) and infrequent ($f_{pos} = 0.1\%$) positive selection. Distribution of (a) nucleotide diversity, (b) Tajima's *D*, (c) $r^2$, and (d) divergence per site across the 465 exons, for each of the three windows separately: functional, linked, and less linked intergenic regions. The best model is depicted in red, the best model with positive selection in blue, and their overlap in purple. The distribution of the empirical data is shown in the white distributions.

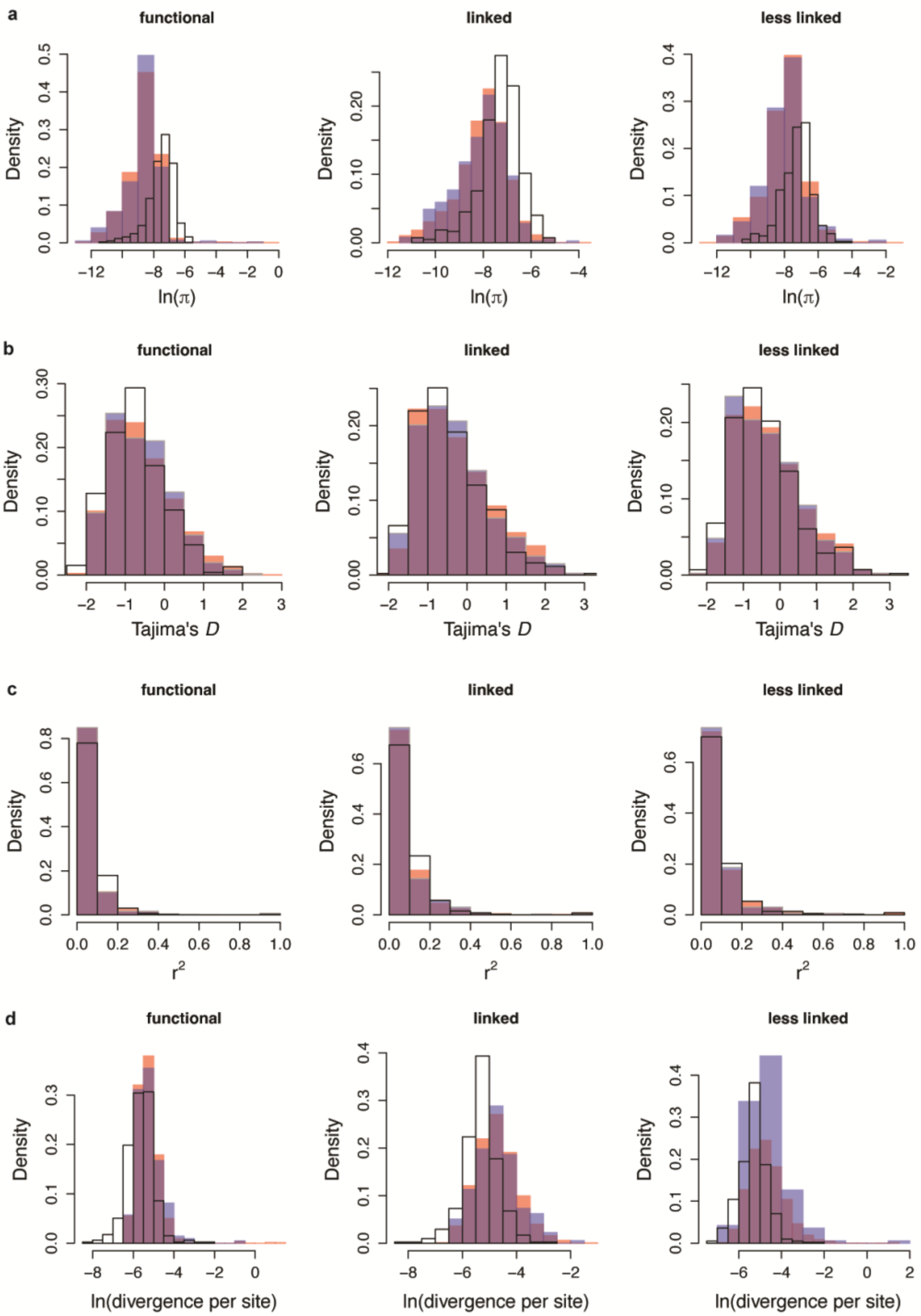


**Figure S18**: Fit of the estimated best model to the empirical data in the presence of weak ($E[2N_e s_b] = 10$) and moderately frequent ($f_{pos} = 1\%$) positive selection. Distribution of (a) nucleotide diversity, (b) Tajima's *D*, (c) $r^2$, and (d) divergence per site across the 465 exons, for each of the three windows separately: functional, linked, and less linked intergenic regions. The best model is depicted in red, the best model with positive selection in blue, and their overlap in purple. The distribution of the empirical data is shown in the white distributions.

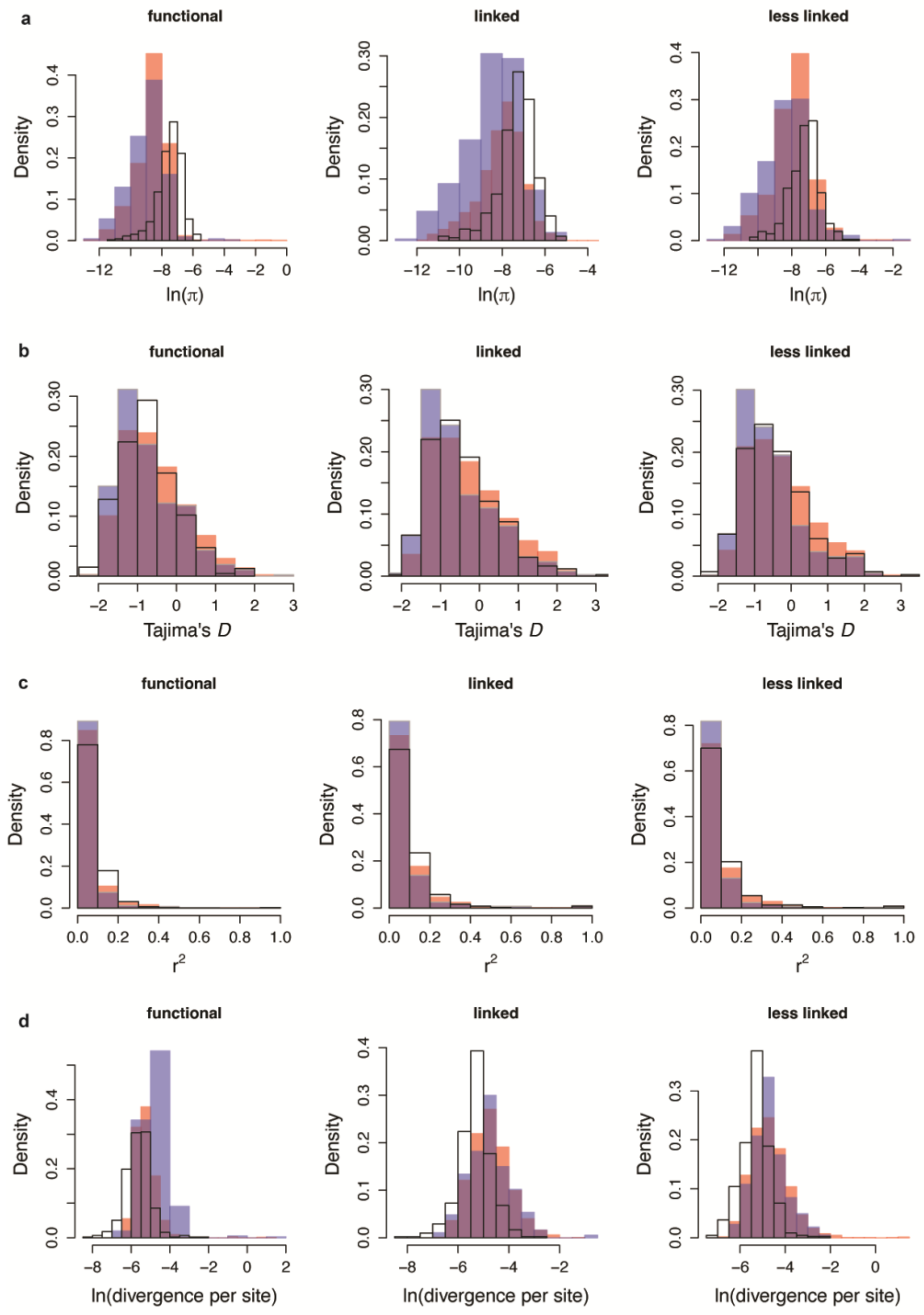


**Figure S19**: Fit of the estimated best model to the empirical data in the presence of moderately strong ($E[2N_e s_b] = 100$) and moderately frequent ($f_{pos} = 1\%$) positive selection. Distribution of (a) nucleotide diversity, (b) Tajima's *D*, (c) $r^2$, and (d) divergence per site across the 465 exons, for each of the three windows separately: functional, linked, and less linked intergenic regions. The best model is depicted in red, the best model with positive selection in blue, and their overlap in purple. The distribution of the empirical data is shown in the white distributions.

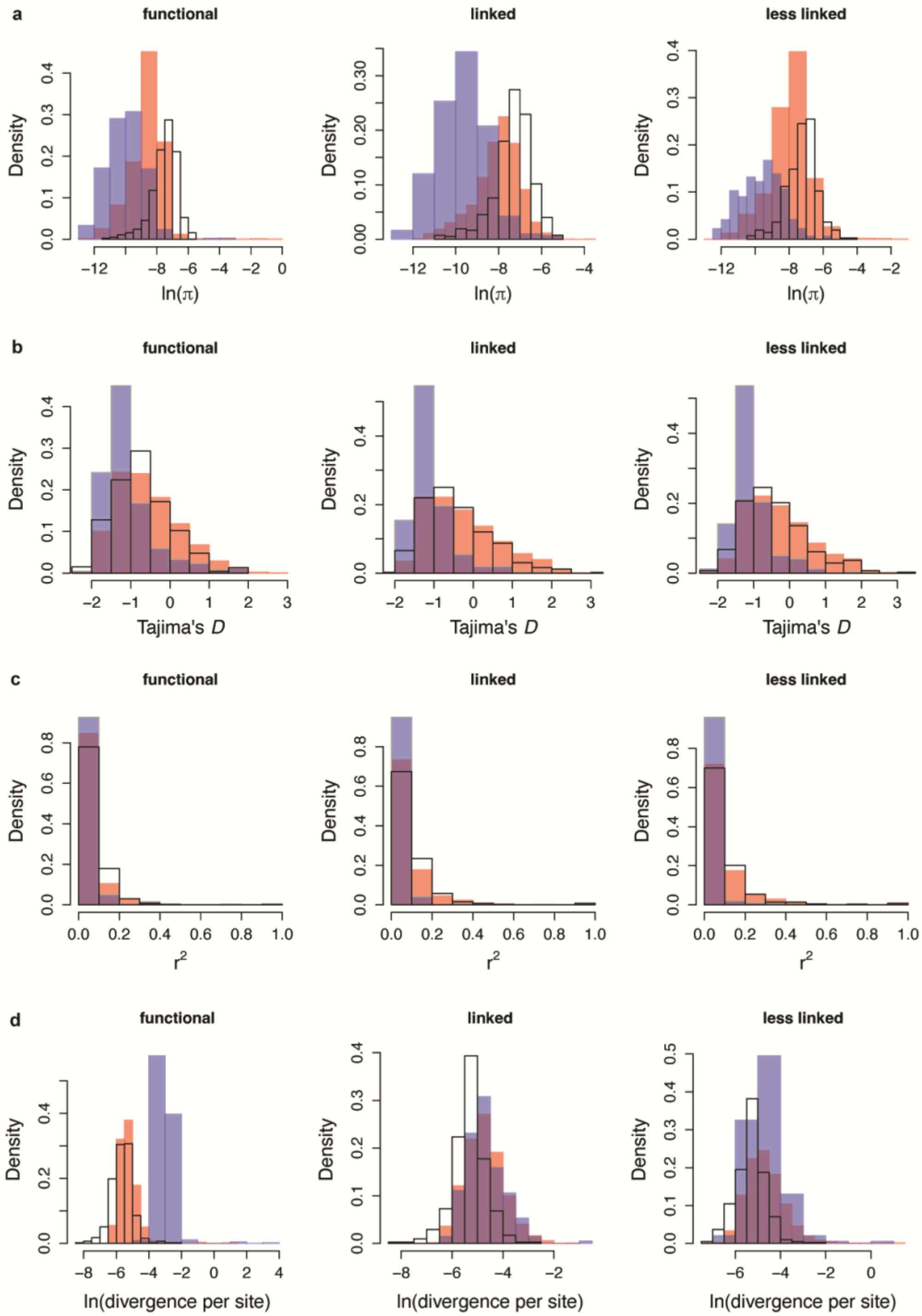


**Figure S20**: Fit of the estimated best model to the empirical data in the presence of strong ($E[2N_e s_b] = 1000$) and moderately frequent ($f_{pos} = 1\%$) positive selection. Distribution of (a) nucleotide diversity, (b) Tajima's *D*, (c) $r^2$, and (d) divergence per site across the 465 exons, for each of the three windows separately: functional, linked, and less linked intergenic regions. The best model is depicted in red, the best model with positive selection in blue, and their overlap in purple. The distribution of the empirical data is shown in the white distributions.

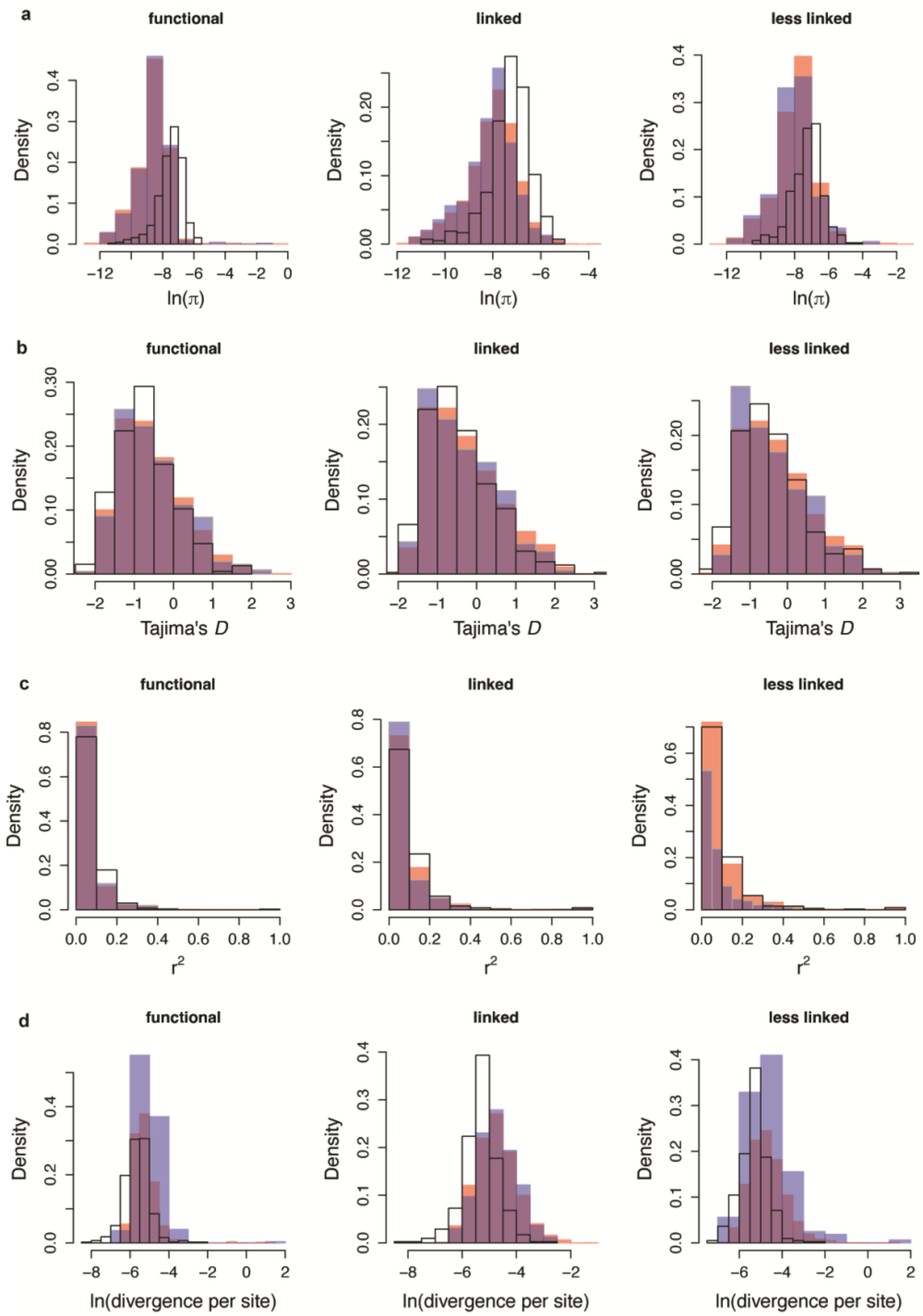


**Figure S21**: Fit of the estimated best model to the empirical data in the presence of weak ($E[2N_e s_b] = 10$) and frequent ($f_{pos} = 5\%$) positive selection. Distribution of (a) nucleotide diversity, (b) Tajima's *D*, (c) $r^2$, and (d) divergence per site across the 465 exons, for each of the three windows separately: functional, linked, and less linked intergenic regions. The best model is depicted in red, the best model with positive selection in blue, and their overlap in purple. The distribution of the empirical data is shown in the white distributions.

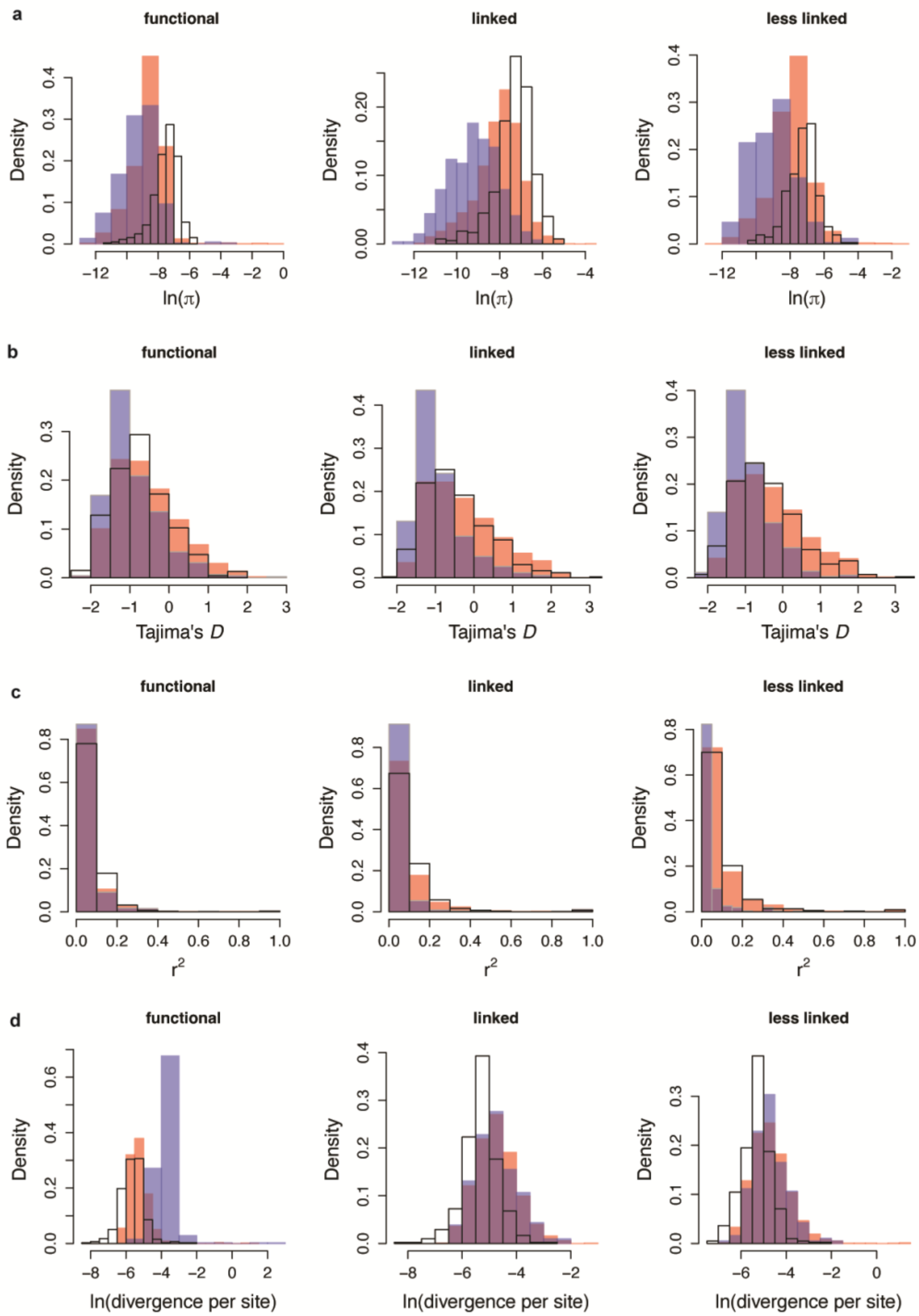


**Figure S22**: Fit of the estimated best model to the empirical data in the presence of moderately strong ($E[2N_e s_b] = 100$) and frequent ($f_{pos} = 5\%$) positive selection. Distribution of (a) nucleotide diversity, (b) Tajima's *D*, (c) $r^2$, and (d) divergence per site across the 465 exons, for each of the three windows separately – functional, linked, and less linked intergenic regions. The best model is depicted in red, the best model with positive selection in blue, and their overlap in purple. The distribution of the empirical data is shown in the white distributions.

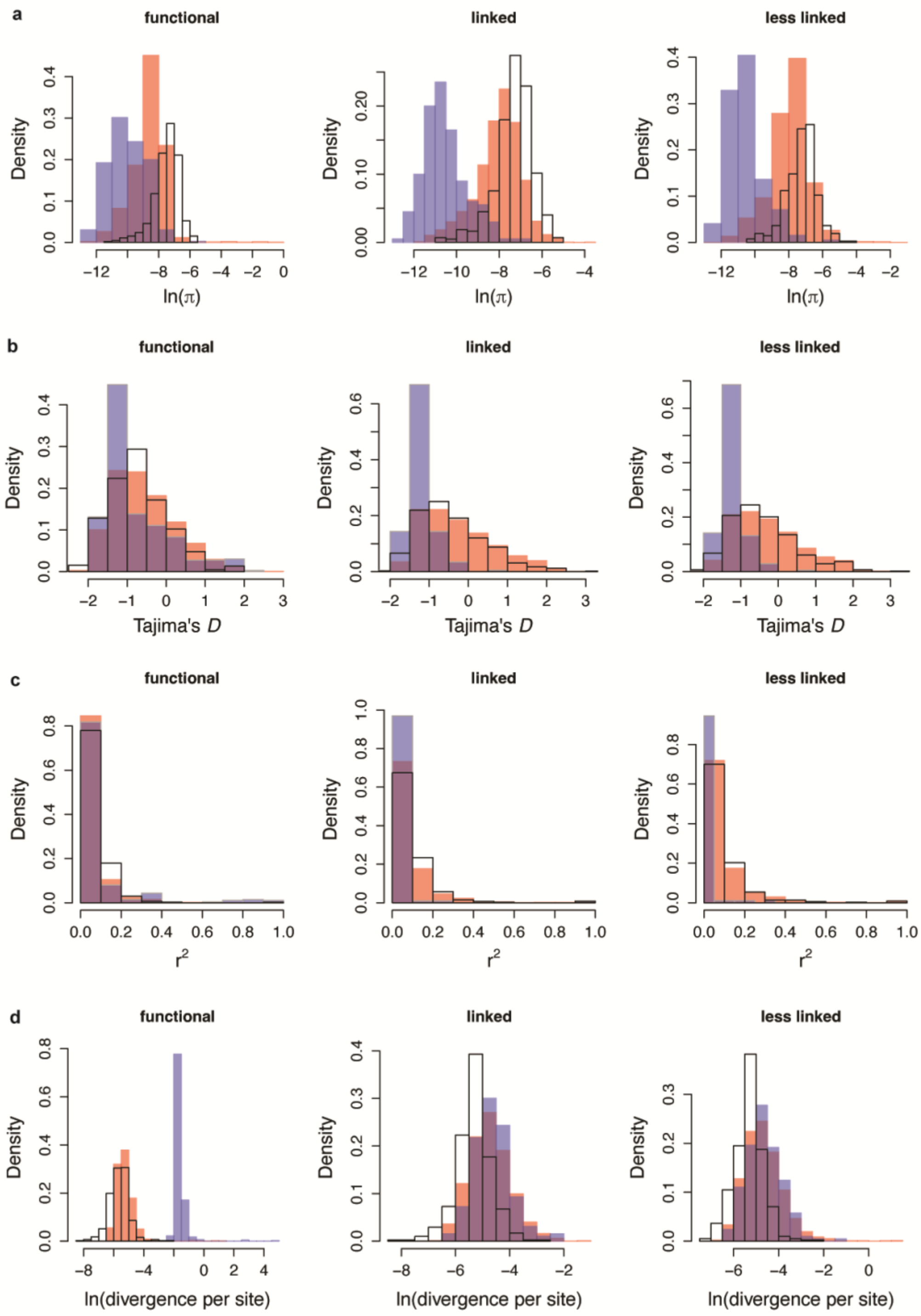


**Figure S23**: Fit of the estimated best model to the empirical data in the presence of strong ($E[2N_e s_b] = 1000$) and frequent ($f_{pos} = 5\%$) positive selection. Distribution of (a) nucleotide diversity, (b) Tajima's *D*, (c) $r^2$, and (d) divergence per site across the 465 exons, for each of the three windows separately: functional, linked, and less linked intergenic regions. The best model is depicted in red, the best model with positive selection in blue, and their overlap in purple. The distribution of the empirical data is shown in the white distributions.

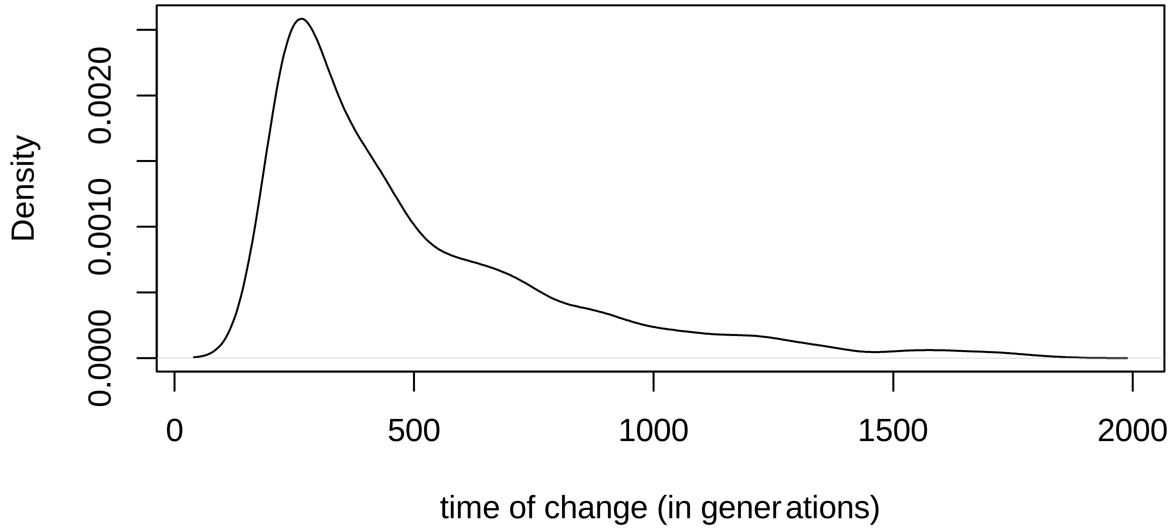


**Figure S24**: Prior distribution of the time of change utilized by our ABC method.

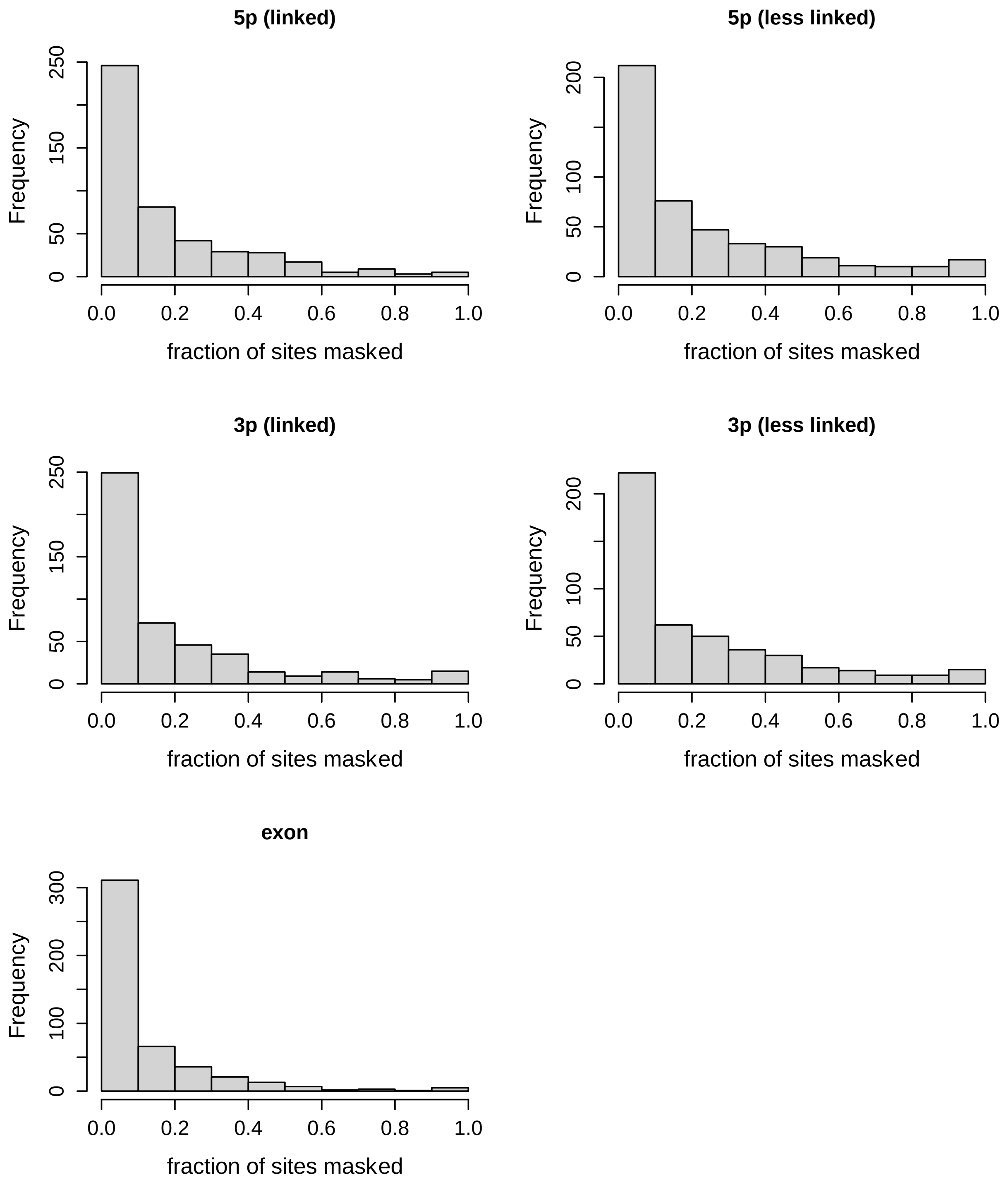


**Figure S25**: Distribution of the proportion of sites masked (or filtered) across the 465 exons and their respective intergenic regions in the YRI 1000 Genomes Phase 3 data.
